# Playing with FiRE: A genome resolved view of the soil microbiome responses to high severity forest wildfire

**DOI:** 10.1101/2021.08.17.456416

**Authors:** Amelia R. Nelson, Adrienne B. Narrowe, Charles C. Rhoades, Timothy S. Fegel, Rebecca A. Daly, Holly K. Roth, Rosalie K. Chu, Kaela K. Amundson, Sara E. Geonczy, Joanne B. Emerson, Robert B. Young, Andrei S. Steindorff, Stephen J. Mondo, Igor V. Grigoriev, Asaf Salamov, Thomas Borch, Michael J. Wilkins

**Affiliations:** Department of Soil and Crop Sciences, Colorado State University, Fort Collins, CO, USA; Rocky Mountain Research Station, U.S. Forest Service, Fort Collins, CO, USA; Department of Chemistry, Colorado State University, Fort Collins, CO, USA; Environmental Molecular Sciences Laboratory, Pacific Northwest National Laboratory, Richland, WA, USA; Department of Plant Pathology, University of California, Davis, CA, USA; Chemical Analysis & Instrumentation Laboratory, New Mexico State University, Las Cruces, NM, USA; Department of Energy Joint Genome Institute, Lawrence Berkeley National Laboratory, Berkeley, CA, USA; Department of Agricultural Biology, Colorado State University, Fort Collins, CO, USA; Department of Plant and Microbial Biology, University of California Berkeley, Berkeley, CA, USA; Department of Civil and Environmental Engineering, Colorado State University, Fort Collins, CO, USA

## Abstract

Warming climate has increased the frequency and size of high severity wildfires in the western United States, with deleterious impacts on forest ecosystem resilience. Although forest soil microbiomes provide a myriad of ecosystem functions, little is known regarding the impact of high severity fire on microbially-mediated processes. Here, we characterized functional shifts in the soil microbiome (bacterial, fungal, and viral) across wildfire burn severity gradients one year post-fire in coniferous forests (Colorado and Wyoming, USA). We generated the **Fi**re **R**esponding **E**cogenomic database (FiRE-db), consisting of 637 metagenome-assembled bacterial genomes, 2490 viral populations, and 2 fungal genomes complemented by 12 metatranscriptomes from soils affected by low and high-severity, and complementary marker gene sequencing and metabolomics data. Actinobacteria dominated the fraction of enriched and active taxa across burned soils. Taxa within surficial soils impacted by high severity wildfire exhibited traits including heat resistance, sporulation and fast growth that enhanced post-fire survival. Carbon cycling within this system was predicted to be influenced by microbial processing of pyrogenic compounds and turnover of dominant bacterial community members by abundant viruses. These genome-resolved analyses across trophic levels reveal the complexity of post-fire soil microbiome activity and offer opportunities for restoration strategies that specifically target these communities.

## Introduction

Changes in climate coupled with the effects of long-term fire suppression and shifting land use patterns have increased the frequency, severity, and season length of wildfires in the western US^1–4^. In 2020 and 2021, much of the western US experienced severe wildfires of record-breaking extent^2^. High severity wildfires cause increased erosion^5^, elevated soil carbon (C) and nitrogen (N) losses^6^, and subsequent nutrient and sediment export in stream water^7,8^, so the increased occurrence of severe wildfires may have important consequences for both terrestrial and aquatic ecosystems. Shifting wildfire patterns have also been linked to slow post-fire revegetation and tree seedling recruitment^9^ and thus delayed watershed recovery^10–12^ in western US forests. Although ecosystem recovery from severe wildfires is closely linked to belowground biological processes, little is known about the impact of high severity fire on soil microbiome function.

The soil microbiome regulates soil organic matter (SOM) decomposition and stabilization^13^, soil nutrient dynamics^14^, and rhizosphere function^15^. During severe wildfires, the soil microbiome can be impacted directly due to heating killing heat-sensitive microbial taxa or indirectly via lasting changes in soil chemistry (e.g., pH, organic matter structure) that continue to influence soil microbiome assembly^16^. Wildfires reduce soil microbial community diversity in numerous ecosystems^17–20^ and such changes are likely to influence and potentially inhibit post-fire plant recovery^21,22^.

Although post-fire shifts in soil microbiome composition are relatively well characterized across ecosystems^16,18,19,23^, the impacts of fire on microbiome metabolic function and microbially-mediated biogeochemical processes are not. To date, the vast majority of soil microbiome studies following wildfire have measured ‘*who is there?*’, rather than focusing on how these compositional shifts affect microbial metabolic functions. While several studies have noted post-fire reductions in genes associated with N-cycling, carbohydrate metabolism, and methanogenesis^24,25^, critical knowledge gaps still exist regarding variability in microbiome responses among and within wildfires due to differences in fire severity (i.e., the degree of vegetation and organic soil horizon combustion). Such information is critical to understanding potential shifts in post-fire ecosystem resilience and to guiding restoration approaches and omics tools have the potential to address this key knowledge gap.

Here, we studied burn severity gradients in two recent forest wildfires in Colorado and Wyoming (USA). Soils were interrogated using a multi-omics approach to characterize how fire severity influences C composition and the intimately connected soil bacterial, fungal, and viral communities. We hypothesized that higher severity wildfire would result in an increasingly altered soil microbiome and that soil microbiomes colonizing burned soils would encode functional traits (e.g., the capacity to utilize fire-altered substrates and rapidly recolonize vacant soil niches) that favor the persistence of specific microbial taxa. Our analysis advances the understanding of specific links between the soil microbiome and post-fire biogeochemical processes associated with forest recovery and provides information critical to land and watershed managers tasked with maintaining the desired ecosystem conditions and the sustained supply of ecosystem services.

## Methods

### Field campaign

Sampling was conducted in lodgepole pine (*Pinus contorta*) forests burned by Badger Creek (8215 ha) and Ryan (11567 ha) fires during 2018 in the Medicine Bow National Forest. Four candidate burn severity gradients were selected based on US Forest Service, Burned Area Emergency Response program (BAER) remotely sensed imagery and maps, and subsequently field validated^26,27^. Aspect, slope, and elevation were recorded at each sampling plot. Each gradient comprised low, moderate, and high severity sites and an unburned control. Low, moderate, and high severity sites had >85%, 20-85%, and <20% surficial organic matter cover, respectively^26^. Samples were collected on August 16 and 19 of 2019, approximately one year following containment of both fires. At each sampling site, a 3 m x 5 m sampling grid with six m^2^ subplots was laid out perpendicular to the dominant slope (**Figure S1**). Subsamples of the organic soil horizon (i.e., litter and duff; O-horizon) and upper mineral soil horizon (0-5 cm; A-horizon) were collected with a sterilized trowel in each subplot for DNA and RNA extractions and subsequent microbial analyses. In three subplots, additional material was collected for chemical analyses. Samples for RNA analyses were immediately flash-frozen using an ethanol-dry ice bath and subsequently placed on ice to remain frozen in the field. Samples for DNA extractions and chemical analyses were immediately placed on ice and all samples were transported to the laboratory at Colorado State University (CSU). Soils for DNA and RNA extractions were stored at -80°C in the laboratory until processing. A total of 176 soil samples were collected (**Supplementary data 1**).

### Soil chemistry

We measured inorganic forms of soil N (NO_3_–N and NH_4_–N) in both organic and upper mineral soils. Ten-gram subsamples were extracted with 50 mL of 2 M KCl within 24 h of sampling and analyzed for NO_3_–N and NH_4_–N by colorimetric spectrophotometry^28^ (Lachat Company, Loveland, CO). A second subsample was oven dried at 105°C for 24 h to determine soil moisture content. (**Supplementary data 1**).

### High-resolution carbon analyses: FTICR-MS

Water extractions were completed on a subset of 47 samples from the Ryan Fire for high-resolution C analyses using Fourier Transform Ion Cyclotron Resonance Mass Spectrometry (FTICR-MS) to analyze dissolved organic matter (DOM) pools. Briefly, 100 mL of milliQ water was added to 50 g of sample in an acid-washed and combusted 250 mL Erlenmeyer flask. These were placed on a shaker table for 10 hours at 170 rpm. Following shaking, liquid was poured off into a 50 mL centrifuge tube and centrifuged for 10 min at 7500 g and supernatant was filtered through a polypropylene 0.2 μm filter (polypropylene material). The extracts were acidified to pH 2 and additionally pre-treated with solid-phase extractions using Agilent Bond Elut-PPL 3 mL columns and diluted to 50 ppm (Agilent Technologies, DE, USA) following standard lab protocol^29^. A 12 Tesla (12T) Bruker SolariX FTICR-MS located at the Environmental Molecular Sciences Laboratory in Richland, WA, USA was used to collect DOM high-resolution mass spectra from each DOM sample. Samples were directly injected into the instrument using a custom automated direction infusion cart that performed two offline blanks between each sample and using an Apollo II electrospray ionization (ESI) source in negative ion mode with an applied voltage of -4.2kV. Ion accumulation time was optimized between 50 and 80 ms. One hundred and forty-four transients were co-added into a 4MWord time domain (transient length of 1.1 s) with a spectral mass window of m/z 100-900, yielding a resolution at m/z 400. Spectra were internally recalibrated in the mass domain using homologous series separated by 14 Da (CH2 groups). The mass measurement accuracy was typically within 1 ppm for singly charged ions across a broad m/z range (100 m/z - 900 m/z). Bruker Daltonics DataAnalysis (version 4.2) was used to convert mass spectra to a list of m/z values by applying the FTMS peak picking module with a signal-to-noise ratio (S/N) threshold set to 7 and absolute intensity threshold to the default value of 100. Chemical formulae were assigned with Formularity^30^ based on mass measurement error < 0.5 ppm, taking into consideration the presence of C, H, O, N, S and P and excluding other elements. This in-house software was also used to align peaks with a 0.5 ppm threshold. The R package ftmsRanalysis^31,32^ was then used to remove peaks that either were outside the desired m/z range (200 m/z – 900 m/z) or had an isotopic signature, calculate nominal oxidation state of carbon (NOSC), and assign Van Krevelen compound classes. Raw FTICR-MS data is provided in archive (doi:10.5281/zenodo.5182305).

Kendrick mass defect (KMD) analysis and plots were employed to identify potential increasing polyaromaticity across the burn severity gradient. The KMD analysis was done using the C_4_H_2_ base unit (50 amu) to represent the addition of benzene to a separate molecular benzene. The mass of each identified ion (M) was converted to its Kendrick mass (KM):

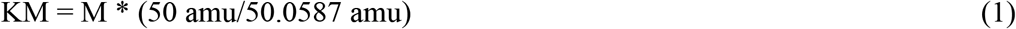

with 50 amu being the nominal mass of C_4_H_2_ and 50.0587 being the exact mass of C_4_H_2_. The final KMD was obtained by subtracting the KM from the nominal KM, which is the initial ion mass rounded to the nearest integer. Series were identified as 2 or more formulas with the same KMD and a nominal Kendrick mass (NKM) differing by the C_4_H_2_ base unit (50 g/mol). Series were retained if they were present across all four burn severity conditions (control, low, moderate, and high), resulting in 64 total series in the final analysis.

### DNA extraction, 16S rRNA gene and ITS amplicon sequencing

Total DNA was extracted from soil samples using the Zymobiomics Quick-DNA Fecal/Soil Microbe Kits (Zymo Research, CA, USA). 16S rRNA genes in extracted DNA were amplified and sequenced at Argonne National Laboratory on the Illumina MiSeq using 251-bp paired-end reads and the Earth Microbiome Project primers 515F/806R^33^, which targets the V4 region of the 16S rRNA gene. To characterize fungal community composition, the DNA was also PCR amplified targeting the first nuclear ribosomal internal transcribed spacer region (ITS) using the primers (ITS1f/ITS2) and sequenced on the Illumina MiSeq platform at the University of Colorado BioFrontiers Institute Next-Gen Sequencing Core Facility using 251-bp paired-end reads.

For taxonomic assignment, we used the SILVA^34^ (release 132) and UNITE^35^ (v8.3) databases for bacteria and fungi, respectively. All sets of reads were clustered into amplicon sequence variant (ASV) classifications using the QIIME2 pipeline^36^ (release 2018.11). 16S rRNA gene and ITS amplicon sequencing data discussed here is available at NCBI under BioProject #PRJNA682830. Ecological guilds were assigned to fungal ASVs using FUNGuild^37^ (v1.2). Guilds were summarized into ‘Saprotroph’, ‘Ectomycorrhizal’, ‘Endophyte’, ‘Epiphyte’, and ‘Arbuscular mycorrhizal’, and were only retained if the confidence ranking was ‘probable’ or ‘highly probable’. If a taxon was classified as multiple guilds that were not consistent in the main guilds listed above (e.g., saprotroph), it was not used for downstream analysis. For example, an ASV classified as ‘Animal Pathogen-Endophyte-Wood Saptrotroph’ would not be retained but an ASV classified as ‘Wood Saprotroph-Soil Saprotroph’ would be retained and renamed ‘Saprotroph’.

To characterize how microbial populations differed across burn severities and soil horizons, we used the R^38^ vegan^39^ (v2.5-7) and phyloseq^40^ (v1.28.0) packages. Nonmetric multidimensional scaling (NMDS) was used to examine broad differences between microbial communities. Analyses of similarity (vegan::anosim) was additionally utilized to statistically test the magnitude of dissimilarity between microbial communities from the different burn severity conditions and soil horizons. Mean species diversity of each sample (alpha diversity) was calculated based on species abundance, evenness, or phylogenetic relationships using Shannon’s Diversity Index (H), Faith’s Phylogenetic Diversity (pd), and Pielou’s Evenness (J). Linear discriminant analysis (LDA) with a score threshold of 2.0 was used to determine ASVs discriminant for unburned or burned soil^41^.

### Metagenomic assembly and binning

A subset of 12 Ryan Fire samples from a single transect representing low and high severity burn from organic and mineral horizon soils was selected for metagenomic sequencing to analyze changes in microbial community functional potential (n=3 per condition). ANOSIM analyses of 16S rRNA gene data confirmed that the soil microbial communities were not significantly different between the Ryan and Badger Creek sites (R = 0.09, p<0.05), so we focused on the Ryan Fire as a representative site. The different conditions are hereafter referred to as ‘Low O’ (low severity organic horizon), ‘High O’ (high severity organic horizon), ‘Low A’ (low severity mineral horizon), and ‘High A’ (high severity mineral horizon). Libraries were prepared using the Tecan Ovation Ultralow System V2 and were sequenced on the NovaSEQ6000 platform on a S4 flow cell using 151-bp paired-end reads at Genomics Shared Resource, Colorado Cancer Center, Denver, CO, USA. Sequencing adapter sequences were removed from raw reads using BBduk (https://jgi.doe.gov/data-and-tools/bbtools/bb-tools-user-guide/bbduk-guide/) and reads were trimmed with Sickle^42^ (v1.33). For each sample, trimmed reads were assembled into contiguous sequences (contigs) using the de novo de Bruijn assembler MEGAHIT v1.2.9 using kmers^43^ (minimum kmer of 27, maximum kmer of 127 with step of 10). Assembled contigs greater than 2500bp were binned using MetaBAT2 with default parameters^44^ (v2.12). Metagenome-assembled genome (MAG) quality was estimated using checkM^45^ (v1.1.2) and taxonomy was assigned using GTDB-Tk^46^ (R05-RS95, v1.3.0). MAGs from all samples were combined and dereplicated using dRep^47^ (default parameters, v2.2.3) to create a non-redundant MAG database. Low quality MAGs (<50% completion and >10% contamination) were excluded from further analysis^48^. Reads from all samples were mapped to the dereplicated bin database using BBMap with default parameters (version 38.70, https://sourceforge.net/projects/bbmap/). Per-contig coverage across each sample was calculated using CoverM contig (v 0.3.2) (https://github.com/wwood/CoverM) with the ‘Trimmed Mean’ method, retaining only those mappings with minimum percent identity of 95% and minimum alignment length of 75%. Coverages were scaled based on library size and scaled per-contig coverages were used to calculate the mean per-bin coverage and relative abundance in each sample (**Supplementary data 2**). The quality metrics and taxonomy of the subsequent 637 medium- and high-quality MAGs discussed here are included in the supplementary material (**Supplementary data 3**) and are deposited at NCBI (BioProject ID PRJNA682830). Maximum cell doubling times were calculated from codon usage bias (CUB) patterns in each MAG using gRodon^49^.

### Metagenome-assembled genome annotation

Eukaryotic MAGs were annotated using the JGI Annotation Pipeline analyzed with complementary metatranscriptomics assemblies^50^ (RnaSPAdes, v3.13.0) and are deposited on MycoCosm^51^ (https://mycocosm.jgi.doe.gov). Bacterial MAGs were annotated using DRAM^52^ (v1.0). In addition to the DRAM annotations, we used HMMER^53^ against Kofamscan HMMs^54^ to identify genes for catechol and protocatechuate meta- and ortho-cleavage, naphthalene transformations, and inorganic N cycling (**Supplementary data 4**). A metabolic pathway within a MAG is considered complete if it is ≥50% complete because MAGs are commonly not 100% complete.

### Metatranscriptomics

Total RNA was extracted from the subset of 12 samples utilized for metagenomics using the Zymobiomics DNA/RNA Mini Kit (Zymo Research, CA, USA) and RNA was cleaned, DNase treated, and concentrated using the Zymobiomics RNA Clean & Concentrator Kit (Zymo Research, CA, USA). Ribosomal RNA was removed from total RNA and libraries were prepared using the Takara SMARTer Stranded Total RNA-Seq Kit v2 (Takara Bio Inc, Shiga, Japan). Samples were sequenced on the NovaSEQ6000 platform on a S4 flow cell using 151-bp paired-end reads at Genomics Shared Resource, Colorado Cancer Center, Denver, CO, USA. Adapter sequences were removed from raw reads using Bbduk (https://jgi.doe.gov/data-and-tools/bbtools/bb-tools-user-guide/bbduk-guide/) and sequences were trimmed with Sickle v1.33^42^. Trimmed reads were mapped to metagenome assemblies using BBMap (parameters: ambiguous=random, idfilter=0.95; v38.70). Mappings were filtered to 95% identity and counts were generated using HTSeq^55^. For broad analysis of differential expression, the dataset was filtered to only transcripts which were successfully annotated by DRAM (n=146,895) and DESeq2^56^ was used to identify transcripts that were differentially expressed in either burn severity within soil horizons (e.g., high vs. low in organic horizon soils and vice versa). The same analysis was also run on the combined HMM output described above (1,189 total transcripts). We normalized our dataset by calculating the gene length corrected trimmed mean of M values^57^ (geTMM) using edgeR^58^ to normalize for library depth and gene length. To identify transcripts that were highly expressed in any given condition, we filtered the data to only transcripts that were in the upper 20% of TMM for 2 of the 3 samples in any one condition (**Table S1**). To compare bacterial and fungal expression data for individual genes, we normalized the number of either fungal or bacterial transcript reads to the gene coverage in each sample to compare the number of transcripts recruited per gene.

### Viruses

Viral contigs were recovered from the metagenomic assemblies using VirSorter2^59^ (v2.2.2) and only contigs >/= 10kb with a VirSorter2 score > 0.5 were retained. Subsequent viral contigs were trimmed using checkV^60^ (v0.4.0) and the final contigs were clustered using the CyVerse app ClusterGenomes (v1.1.3) requiring an average nucleotide identity of 95% or greater over at least 80% of the shortest contig. The final DNA viral metagenome-assembled genomes (vMAG) dataset was manually curated using the checkV, VIRSorter2, and DRAM-v annotation outputs according to standard protocol^61^. RNA vMAGs were also recovered from metatranscriptome assemblies using VIRSorter2^59^ (v2.2.2). The resulting sequences were clustered using ClusterGenomes (v1.1.3) on CyVerse using the aforementioned parameters. To quantify relative abundance of DNA and RNA vMAGs across the 12 samples, we mapped the metagenomic and metatranscriptomic reads to the vMAGs using BBMap with default parameters (v38.70). To determine vMAGs that had reads mapped to at least 75% of the vMAG, we used CoverM (v0.6.0) in contig mode to find vMAGs that passed this 75% threshold (--min-covered-fraction 75). We then used CoverM (v0.6.0) in contig mode to output reads per base and used this to calculate final DNA and RNA vMAG relative abundance in each metagenome and metatranscriptome. vConTACT2 (v0.9.8; CyVerse) was used to determine vMAG taxonomy. Final viral sequences are deposited on NCBI (BioProject ID #PRJNA682830 - BioSamples SAMN20555178, SAMN20555179; **Supplementary data 5**). We used DRAM-v^52^ (v1.2.0) to identify auxiliary metabolic genes (AMGs) within the final viral database^52^ (**Supplementary data 6**).

CRISPR-Cas protospacers were found and extracted from MAG sequences using the CRISPR Recognition Tool^62^ (CRT, minimum of 3 spacers and 4 repeats) in Geneious (v2020.0.3) and CRisprASSembler^63^ with default parameters (v1.0.1). BLASTn was used to compare MAG protospacer sequences with protospacer sequences in vMAGs with matches only retained if they were 100% or contained ≤1 bp mismatch with an e-value ≤1e-5. To identify putative vMAG-MAG linkages, we used an oligonucleotide frequency dissimilarity measure (VirHostMatcher) and retained only linkages with a d_2_^*^ value < 0.25^64^ (**Supplementary data 7**).

## Results & Discussion

### Increasing fire severity drives decreasing diversity and compositional shifts in the soil microbiome

Near surface soils were collected approximately one year post-fire from four burn severity gradient transects (control, low, moderate, and high burn severity) across two distinct wildfires that occurred in 2018 along the Colorado-Wyoming border. Bacterial and fungal communities in all samples were profiled using marker gene analyses, while a subset of twelve samples (soils impacted by either low or high wildfire severity within the Ryan fire) were additionally interrogated with metagenomic and metatranscriptomic sequencing. Bacterial and fungal communities within burned and unburned soils were significantly different (bacterial ANOSIM R = 0.57, p<0.05; fungal ANOSIM R = 0.72, p<0.05) (**Figure S2**). Reflecting observations from prior studies^17,20,65,66^, bacterial communities in burned soils were characterized by lower diversity and were enriched in Actinobacteria and Bacteroidetes relative to unburned controls (**Figure 1**). Specifically, the Actinobacteria genera *Arthrobacter, Modestobacter, Blastococcus, and Actinomadura* had the largest relative abundance increases in burned soils relative to control soils and were discriminant features of these conditions (**Supplementary data 8**). The diversity of fungal communities in surficial soils also decreased with fire (**Figure 2E**) and, similar to previous studies^18,67,68^, shifted from Basidiomycete- to Ascomycete-dominated with the Basidiomycota relative abundance decreasing by ∼58% (**Figure 1**). Discriminant fungal taxa included ASVs from the *Sordariomycetes, Saccharomycetes*, and *Dothideomycetes*, taxa also found in previous fire studies^66,68^ (**Supplementary data 8**).

**Figure 1.**
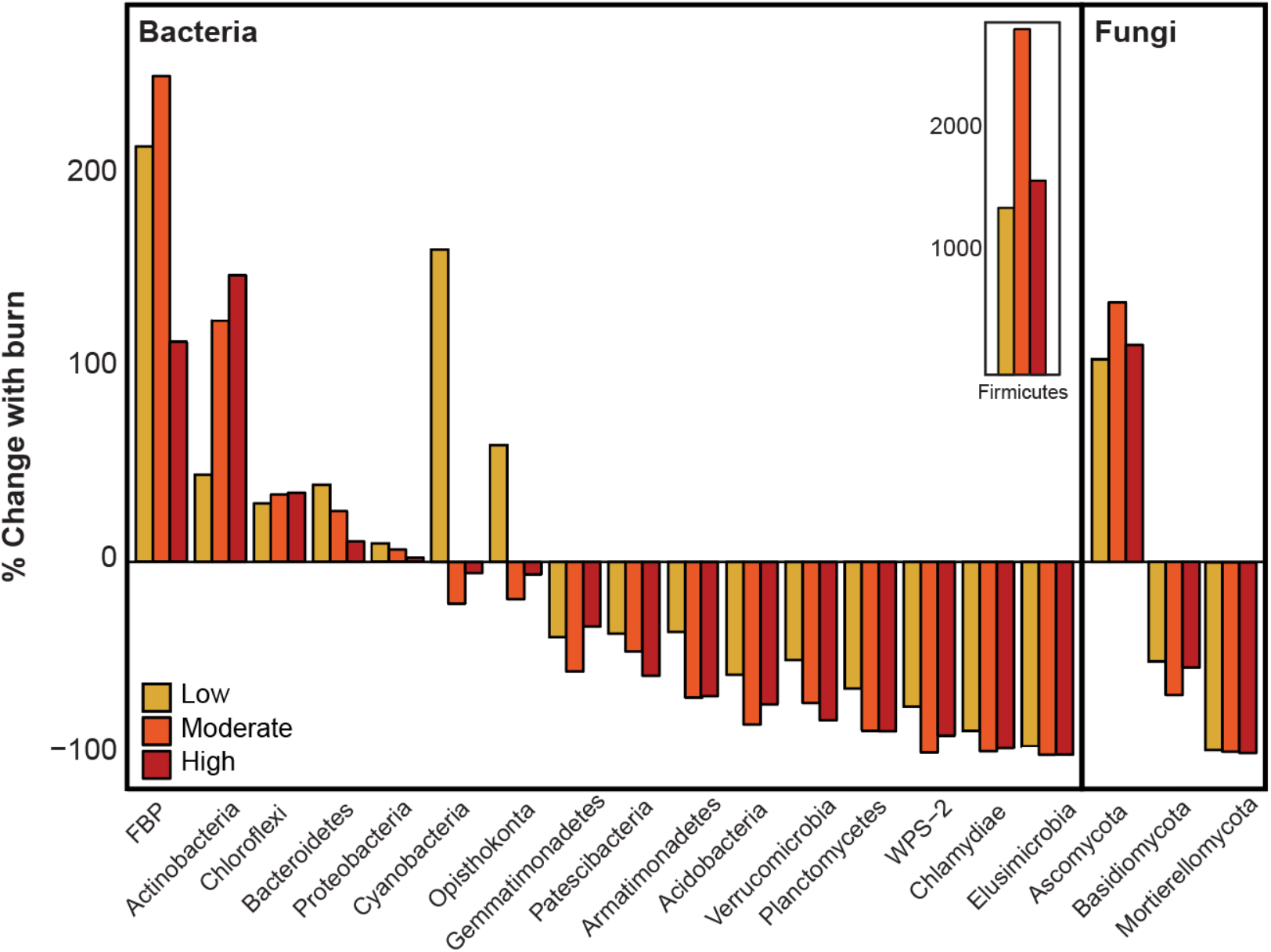
The percent change in relative abundance from control to low, moderate, and high severity in organic soil horizons of each main bacterial and fungal phylum. Phyla with relative abundance less than 0.5% were discarded for this analysis. Note that although the *Firmicutes* have the largest increase with burn (inset) their overall relative abundance is still low.

While shifts in community composition between burned and control conditions were observed in both organic and mineral soil horizons, the surficial organic layers changed more than the deeper mineral horizon soil. Microbial diversity generally decreased with increasing fire severity in the organic horizons, although differences between moderate and high severity were statistically indistinct (**Figure 2**). Similarly, as fungal and bacterial diversity decreased with burn severity, beta dispersion (‘distance to centroid’) calculations revealed increasingly similar microbial community structures (**Figure S3**) with a less complex bacterial and fungal community structure (**Table S2**). These shifts resulted in significant dissimilarity between microbial communities in organic horizons impacted by either low or high severity wildfire (bacterial ANOSIM R=0.15, p<0.05; fungal ANOSIM R=0.25, p<0.05). In contrast, mineral soils that were less impacted by wildfire displayed an opposite effect, with increasing beta dispersion after wildfire that signifies greater bacterial community dissimilarity (**Figure S3**). Stochastic community shifts in deeper soils may follow wildfire, potentially due to spatially heterogeneous changes in soil chemistry and nutrient availability. Together, these data highlight the susceptibility of highly combustible surficial organic soil horizons to wildfire, resulting in less diverse and inter-connected microbial communities. In contrast, mineral soils are likely more insulated from the effects of fire, and therefore the soil microbiome displays a more muted response to wildfire effects.

**Figure 2.**
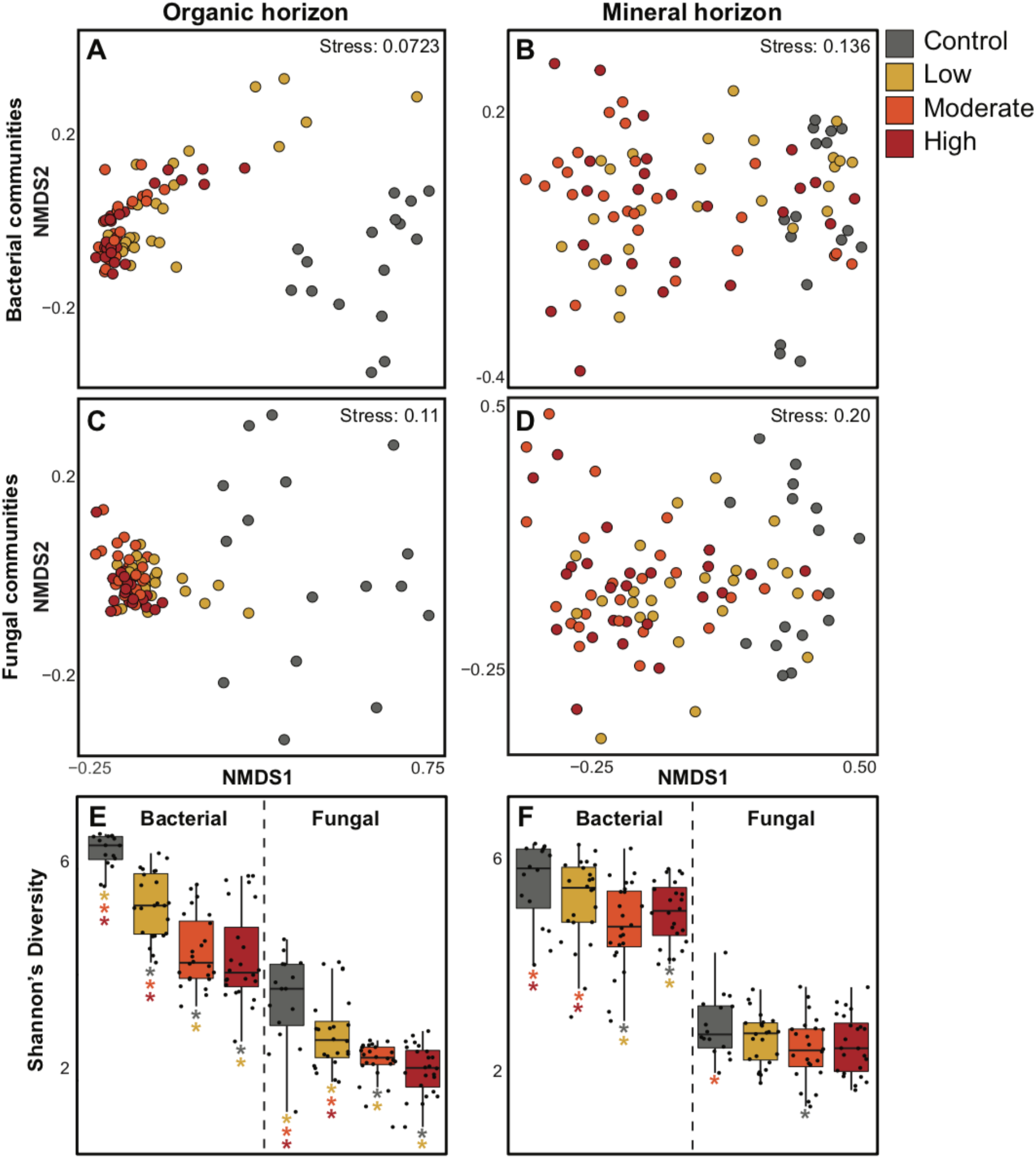
Non-metric multidimensional scaling (NMDS) of organic **(A, C)** and mineral soil horizons **(B, D)** bacterial **(A, B)** and fungal **(C, D)** communities shows increased separation of burned and unburned microbial communities in surficial organic horizon relative to deeper mineral soil communities. Shannon’s diversity (H) calculated from 16S rRNA and ITS gene sequencing in organic **(E)** and mineral soil horizons **(F)** further shows the increased susceptibility of microbiomes in organic horizons to wildfire.

### Development of a unique MAG database from fire-impacted soils

While a myriad of studies have reported changes in microbial community membership in response to wildfire^66,69–71^, the functional implications of these shifts are difficult to infer from marker gene studies. Here we used genome-resolved metagenomics to generate the **Fi**re **R**esponding **E**cogenomic database (FiRE-db), a comprehensive, publicly accessible database of fire-responding bacterial, fungal, and viral genomes from coniferous forest soils (BioProject ID #PRJNA682830). From metagenomics sequencing of 12 burned (low and high severity) forest soil samples we reconstructed 637 medium- and high-quality bacterial metagenome assembled genomes (MAGs) that reflected the majority of dominant taxa observed in complementary 16S rRNA gene analyses (**Figure S4**). This database spans 21 phyla and encompasses 237 genomes from taxa within the Actinobacteria, 167 from the Proteobacteria, 62 from the Bacteroidota, and 52 from the Patescibacteria. Furthermore, we recovered 2 fungal genomes from the Ascomycota, which were affiliated with *Leotiomycetes* and *Coniochaeata lignaria*. We additionally recovered 2,399 DNA and 91 distinct RNA vMAGs from the 12 metagenomes and metatranscriptomes (**Table S5**).

### Actinobacteria respond strongly to high severity wildfire disturbances in surface soils

Reflecting observations from 16S rRNA gene analyses, MAGs affiliated with the Actinobacteria genera *Blastococcus, Mycetocola, SISG01, SCTD01, Nocardiodes,* and *Arthrobacter* were all enriched (relative to control soils) in surface organic soil horizons (O-horizon) impacted by high severity wildfire (hereafter referred to as ‘High O’) that in most instances had been combusted to an ash layer. Because of their enrichment following wildfire, we have focused on 8 featured Actinobacteria MAGs (MAGs RYN_93, RYN_94, RYN_101, RYN_124, RYN_147, RYN_169, RYN_175, RYN_216) for analyses described below. Combined, these MAGs accounted for an average relative abundance of 19% in High O soils and 12% in O-horizon soils impacted by low severity wildfire (‘Low O’ soils). Furthermore, metatranscriptomic analyses indicated that these MAGs were also among the most active in High O samples, accounting for an average of nearly 4% of gene expression across samples. These MAGs were also active in Low O samples, but to a lesser extent (accounting for 1.5% of expression). Together, all 237 Actinobacteria MAGs were responsible for ∼51% of gene expression in the High O soils and ∼42% in Low O samples.

High severity wildfire exerts a pulse disturbance on surficial organic soil horizons by exposing them to high heat. Thus, the taxa that constitute the post-fire microbiome and fill new niches in the soil likely encode traits for thermal resistance. Nearly all the aforementioned Actinobacterial MAGs encoded sporulation genes, indicating that spore formation is likely a key strategy to ensure survival and colonization post-fire (**Figure 3A**). These MAGs also all contained genes encoding heat shock proteins and molecular chaperones which may further facilitate thermal resistance (**Figure 3A**). In the majority of these MAGs, thermal resistance was complimented by genes for mycothiol biosysnthesis, a compound produced by Actinobacteria that aids in oxidative stress tolerance^72^. Finally, MAG RYN_93 encoded *ectB* for synthesizing ectoine, a compatible solute for environmental stress tolerance^73^. In general, MAGs recovered from High O samples also had significantly higher GC content than MAGs from mineral soil layers (e.g., A horizons) impacted by low (‘Low A’) and high (‘High A’) wildfire severity (**Figure S5**). Higher GC content may be another heat resistance trait due to the thermal stability of the GC base pair^74,75^. The heating of organic soil horizons during wildfire lyses heat-sensitive microorganisms, resulting in an influx of labile organic C and N associated with necromass^76^. This likely opens up niche space to fire-resistant heterotrophic taxa and stimulates growth rates of these taxa^77^. Each of the abundant featured Actinobacteria MAGs expressed peptidase genes (88 total genes) in High O samples, of which approximately twenty were differentially expressed (p<0.05) between High O and Low O conditions. These included genes responsible for the degradation of peptidoglycan (component of bacterial cell walls) and chitin (component of fungal cell walls), suggesting that taxa enriched post-fire actively utilize microbial necromass.

**Figure 3.**
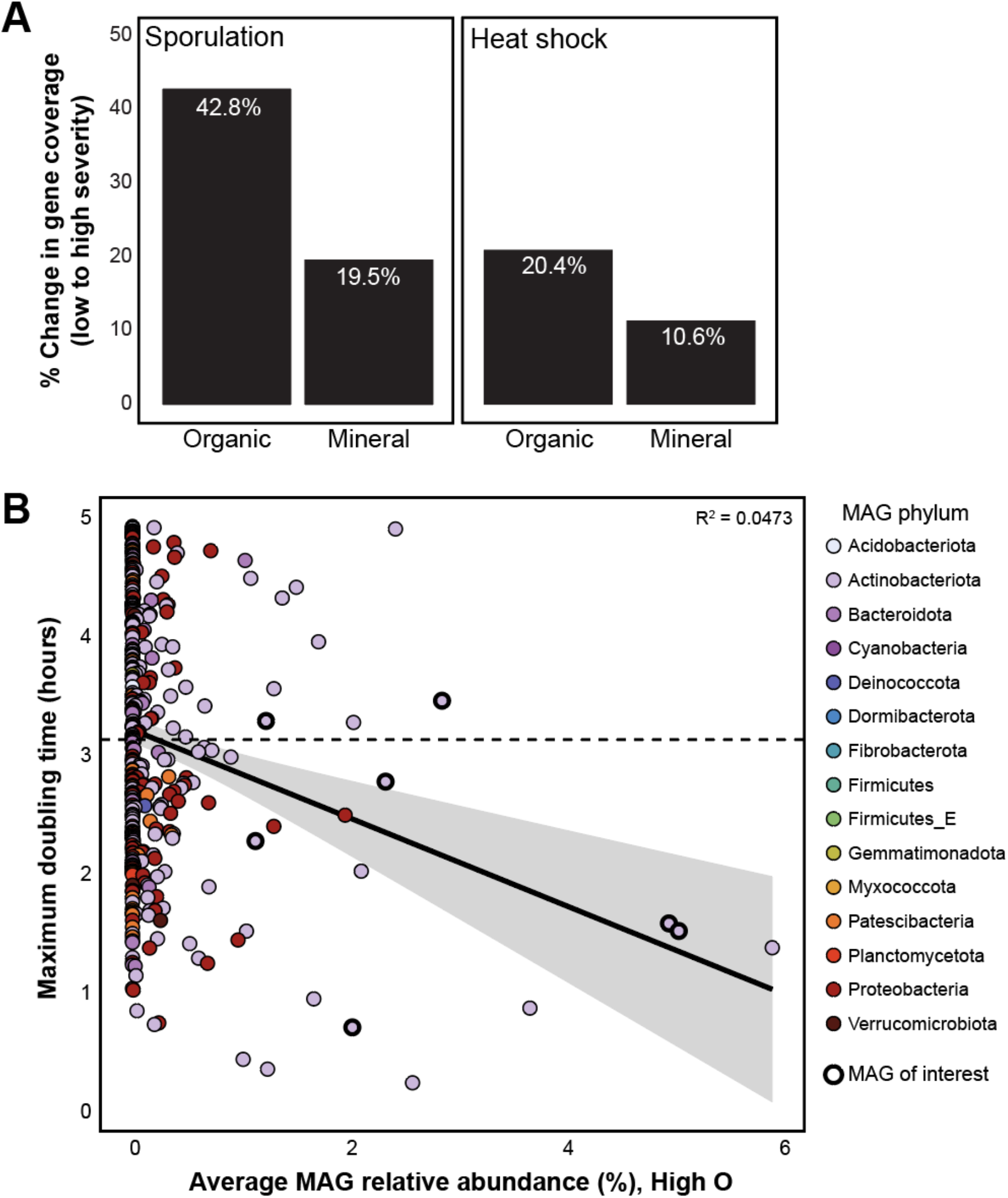
**(A)** Genes encoding for heat resistant traits (sporulation and heat shock) increase in coverage from low to high severity conditions in the organic and mineral soil horizon metagenomes, but to a lesser degree in the mineral horizon soils. **(B)** The MAGs of interest that are enriched and highly active in High O generally have a faster growth rate (lower maximum doubling time, estimated using gRodon^49^) than the average growth rate (indicated here with the dashed line) in our MAG database.

The ability to grow quickly and occupy available niches in the environment is likely a key trait for microorganisms colonizing burned soils^66,78^. We inferred maximum growth rates across our bacterial MAGs to determine whether colonizing taxa encoded the potential for rapid growth in burned soils^49^. Potential growth rates (as maximum cell doubling times) were calculated from patterns of codon usage bias (CUB) in each MAG^79,80^. After removal of all MAGs with doubling time greater than 5 hours due to model inaccuracies at slower growth rates^49^, the average doubling time within our MAG database was 3.16 hours. All but two of the dominant MAGs in High O samples had doubling times faster than the database average (ranging from 0.73 to 3.59 hours) (**Figure 3B)**. These insights suggest that abundant bacteria sampled one year post-wildfire likely occupied niches in the immediate aftermath of wildfire through rapid cell growth. In contrast, these patterns were absent from MAGs from High A soils (**Figure S7**). Emphasizing the importance of fast growth for colonizing severely burned soils, only 19 bacterial MAGs from High O samples had growth rates too slow for the model to accurately estimate (total of 249 MAGs with growth rates >5 hours). To determine whether these same microorganisms were growing rapidly at the time of sampling (one year post-wildfire), we investigated gene expression associated with rapid growth (i.e., ribosomes, central metabolism) through MAG abundance-normalized transcripts. Results suggested diminished growth rates for the dominant High O MAGs at time of sampling, relative to other Actinobacteria MAGs (e.g., *Blastococcus, Nocardiodes*) that accounted for high expression of ribosomal genes and components of the TCA cycle in High O samples. Together, these analyses indicate that the rapid growth rates enabling ‘fast-responders’ to occupy free niche space in soil immediately following wildfire are not maintained once those niches are filled.

### Deeper burned soils are dominated by distinct microbial membership

High A soils hosted a greater diversity of enriched and active MAGs relative to the overlying organic soil horizons. Actinobacteria again contributed strongly to these signals, with the dominant, active MAGs in the mineral horizons affiliated with the families *Streptosporangiaceae, Solirubrobacteraceae, Frankiaceae*, and *Streptomycetaceae* (MAGs RYN_173, RYN_225, RYN_228, RYN_220, RYN_230, RYN_265). Additional highly abundant and active MAGs in High A samples were affiliated with the Eremiobacterota, Dormibacterota, and Proteobacteria phyla (MAGs RYN_132, RYN_309, RYN_342, RYN_347, RYN_607). Together, these MAGs accounted for ∼20% of the High A community composition, and nearly 14% of total MAG gene expression. These MAGs were also active in Low A soils, albeit to a lesser extent (accounting for ∼5% of total expression).

Genes associated with thermal resistance were again common in the dominant High A MAGs. Three of the MAGs discussed here (RYN_220, RYN_225, RYN_342) had metabolic potential to synthesize the environmental stress protectant mycothiol^72^. The second most active MAG in High A samples (RYN_220) was affiliated with the *Streptomyces* which have been shown to form sporogenic structures (aerial hyphae) in response to adverse conditions (i.e., high temperatures, nutrient depletion)^81^. We observed an overall increase in spore-forming gene coverage from Low A to High A conditions (**Figure 2A**) and all MAGs discussed here encoded at least one sporulation gene. Additionally, members of *Streptomyces* are known for their ability to scavenge nutrients and utilize a diverse array of organic substrates^82^, which is evidenced here through expression of abundant and diverse CAZymes that are predicted to target chitin, polyphenolics, and pectin, among other substrates (**Figure S6**).

Associated with the enrichment and activity of *Streptomyces* in High A soils, we observed high expression of genes encoding the biosynthesis of cobalamin (vitamin B12, *cob* genes). Cobalamin production is conserved within a relatively small group of microorganisms – including *Streptomyces* – and can serve as a keystone function within ecosystems^83^. The entire aerobic cobalamin biosynthesis pathway was expressed in Low A and High A samples (**Figure S8**; pathway adapted^83,84^). Here, a *Streptomyces* MAG (RYN_220) was responsible for 12% of MAG gene expression linked to cobalamin biosynthesis in High A soils. In total, the MAGs mentioned above were responsible for nearly 90% of the total cobalamin biosynthesis MAG gene expression in High A samples. These observations contrast directly with High O samples; the general absence of *Streptomyces* in these samples resulted in limited expression of this pathway. Cobalamin biosynthesis gene expression was ∼175% greater in High A samples, and 3 of the aforementioned MAGs (RYN_342, RYN_225, RYN_220, RYN_347) differentially expressed *cobB, cobA/O*, and *btuB* (cobalamin transporter gene) in High A soils relative to High O. In the A-horizon, the increased transcription of genes for cobalamin synthesis is likely a beneficial consequence of wildfire enriching taxa that encode this trait (i.e., *Streptomyces*). Given the noted importance of this cofactor in mediating a range of critical soil microbiome functions, this process could potentially aid in plant reestablishment^85^, enhance ecosystem function across trophic levels^86^, and thus be beneficial for the overall restoration of post-fire landscapes.

### Actinobacteria catalyze degradation of pyrogenic organic matter

During wildfire, SOM may be transformed to increasingly aromatic molecular structures that are commonly considered less labile for microbial utilization^87^. Mass spectrometry DOM analyses found evidence for increasing aromaticity with burn severity in organic horizon soils (**Figure 4**). Low severity wildfire drives an increase in DOM aromaticity (**Figure 4B**) but also an accumulation of other unique compounds likely resulting from incomplete combustion of SOM^88^ (**Figure 4A**). In contrast, the unique compounds formed following moderate and high severity wildfire were constrained to the aromatic region (**Figure 4A**). Reflecting the more insulated, deeper mineral soils, these aromaticity index trends were not identified in DOM released from mineral horizons (**Figure S9**). To estimate lability of these DOM pools, we calculated NOSC which can reveal the potential thermodynamic favorability of a carbon substrate, with higher NOSC values theoretically yielding a lower ΔG _C ox_ (i.e., more favorable) when coupled to the reduction of an electron acceptor^89^. Unique formulas detected in High O samples had significantly higher NOSC values than both Low O and control samples, indicating increasing thermodynamic favorability of DOM following severe wildfire (**Figure 4C**).

**Figure 4.**
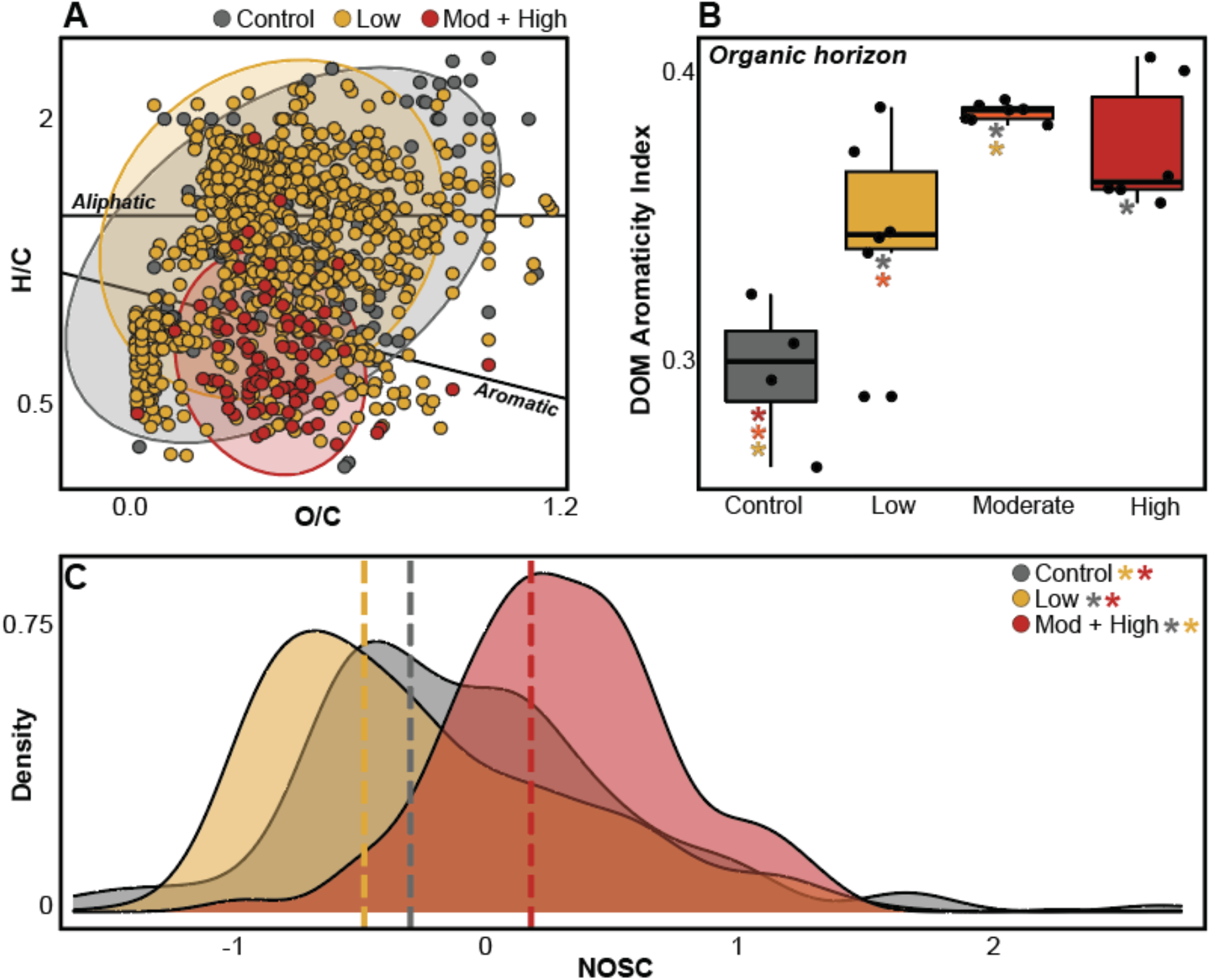
**(A)** Van Krevelen diagram showing unique formulas from organic soil horizons in unburned, low, and moderate and high (combined) organic soil horizons. **(B)** Aromaticity index of DOM pools extracted from organic soil horizons across the burn severity gradient. Colored asterisks indicate significant difference between the two conditions (p<0.05). **(C)** Density plot of unique formula NOSC value distributions between different conditions. Dashed lines show NOSC medians for each condition.

Wildfire severity-driven chemical changes likely mediate the extent of microbial DOM processing. We focused on the ability of community members to process catechol and protocatechuate, the two intermediate products formed during aerobic degradation of diverse aromatic compounds^90^. The genomic potential for these reactions was broadly distributed across burn severities and was dominated by Actinobacteria and Proteobacteria (**Figure 5**); 80 and 226 MAGs encoded at least 50% of the catechol and protocatechuate ortho-cleavage pathways, respectively, including most of the featured High O and High A MAGs (**Figure 5C**). Meta-cleavage pathways were also broadly represented within the MAG database (**Figure S10**). There was additional evidence for activity of both pathways regardless of soil horizon and wildfire severity (**Figure 5A**). In High O samples, the *Arthrobacter* MAG RYN_101 alone was responsible for ∼44% of *catA* (catechol 1,2-dioxygenase) gene expression, and therefore likely plays a key role in catechol degradation. Contrastingly, in High A samples, the *Streptosporangiaceae* MAG RYN_225 was responsible for ∼46% and 23% of the expression of *pcaGH* (protocatechuate 3,4-dioxygenase) and *pcaC* (4-carboxymuconolactone decarboxylase) that catalyzes protocatechuate degradation (**Figure 5C**). However, none of the dominant MAGs across High O and High A samples encoded the entire catechol or protocatechuate ortho-cleavage pathway (**Figure 5C**), indicating that metabolic handoffs between community members are likely important for complete degradation of these compounds.

**Figure 5.**
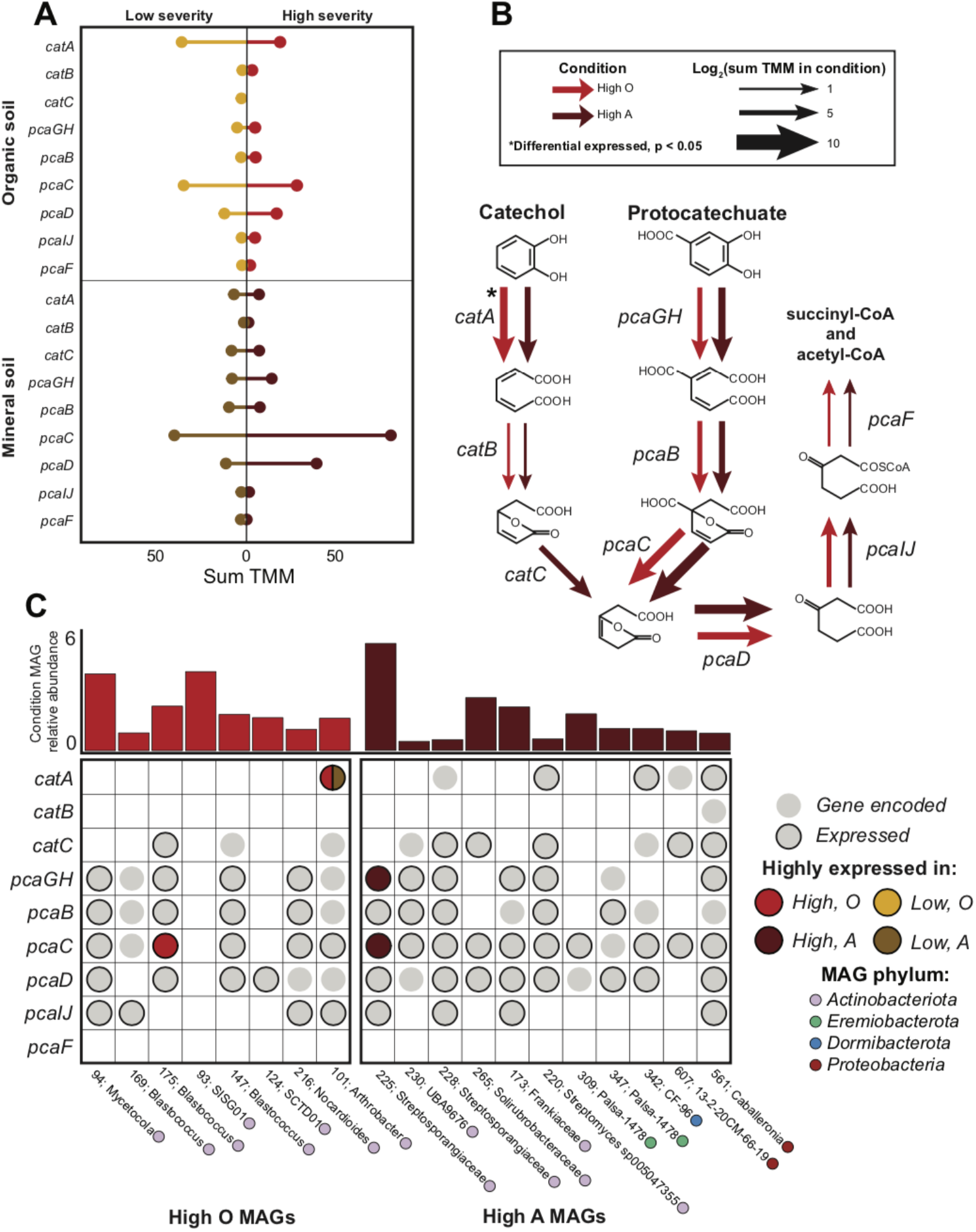
**(A)** The summed TMM of each gene for catechol and protocatechuate ortho-cleavage in each condition. **(B)** The pathway for catechol and protocatechuate ortho-cleavage with arrows indicating the log normalized sum TMM of the gene for high severity organic and mineral soil horizons. Asterisks indicate genes that are differentially expressed in the condition (p<0.05). **(C)** The genomic potential and expression of each gene in the pathway for the MAGs enriched and active in high severity organic and mineral soil samples. The above bar chart shows the featured MAG relative abundance in that condition, colored by featured condition.

### Viral dynamics impact community structure and function of the burned soil microbiome

We recovered 2399 distinct DNA viral populations and 91 distinct RNA viral populations (vMAGs) from the metagenomic and metatranscriptomic assemblies. Highlighting the viral novelty in these ecosystems, 945 of the DNA and RNA vMAGs were previously undescribed (only clustering with other vMAGs from this study) and 92 were given taxonomic assignments, with the majority (n=86) within the *Caudovirales* order (**Table S5**). Both DNA and RNA viral communities mirrored beta diversity trends observed in the bacterial and fungal communities; those in mineral soil horizons were less constrained in multivariate space compared to communities in organic horizons, further highlighting the homogenizing influence of wildfire (**Figure S11**). Additionally, DNA and RNA viral community composition was not significantly different between low- and high-severity impacted soils (ANOSIM R = 0.007 and -0.12, respectively; p > 0.1) but significantly differed between the two soil horizons (ANOSIM R = 0.59 and 0.57, respectively; p<0.05).

Given the importance of viral activity on soil microbiome community composition and function^91,92^, we identified potential virus-host linkages that could offer insights into how viruses target bacteria. Many abundant and active MAGs (n=94) – including 32 from the Actinobacteria– encoded CRISPR-Cas arrays with an average of ∼18 spacers (max 210 spacers; **Supplementary data 3**). Through the matching of protospacers to sequences in vMAGs, we linked 9 vMAGs with 4 bacterial hosts (RYN_115, RYN_242, RYN_436, RYN_542) from the phyla Actinobacteria, Planctomycetota, and Proteobacteria. While each of these four MAGs were active (via transcript expression), the RYN_242 MAG (*Solirubrobacteraceae*) was among the top 3% of active MAGs across all four conditions, suggesting that viruses are targeting active bacteria. We expanded upon potential virus-host linkages using VirHostMatcher^64^ (d_2_^*^ value < 0.25), revealing higher numbers of viral linkages with more abundant host MAGs (**Figure 6**). For example, the MAGs of interest from High O and High A samples had above average numbers of putative viral linkages (average of 196; **Figure 6**). Moreover, there were 198 vMAGs with putative linkages to all 8 featured Actinobacteria MAGs from High O soils, potentially due to conserved nucleotide frequencies. The shared 198 vMAGs made up ∼11% of the viral community in High O samples, again suggesting that the most abundant and active bacteria in burned soils are being actively targeted by abundant phage, potentially impacting soil C cycling via release of labile cellular components following cell lysis. There is also evidence in this system for the “piggyback-the-winner” viral strategy, where lysogenic lifestyles are favored at high microbial abundances and growth rates^93^. Of our 2399 DNA vMAGs, we identified 185 with putative lysogenic lifestyles based on gene annotations for integrase, recombinase, or excisionase genes, 32 of which had nucleotide frequency-based linkages to all the featured Actinobacteria MAGs in High O samples.

**Figure 6.**
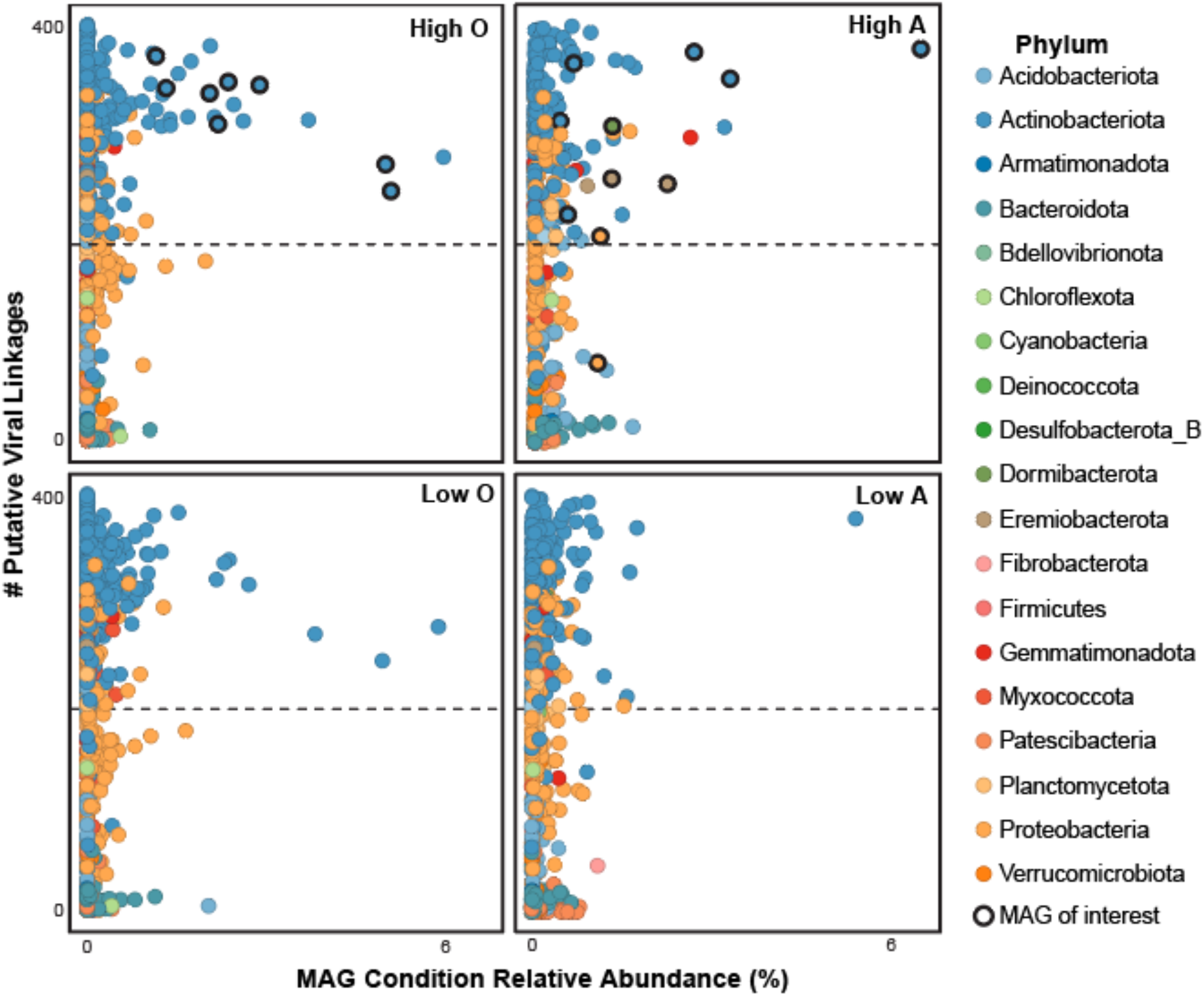
Each MAG’s relative abundance within each respective condition plotted against the number of putative viral linkages identified by VirHostMatcher. Dashed line indicates the database average of 196. Points outlined in bold represent MAGs discussed as important in either High O or High A samples.

To further investigate potential viral roles in C cycling in burned soils, we characterized the putative AMG repertoire of the vMAGs. Viruses utilize AMGs to “hijack” and manipulate the metabolisms of their host; one study in thawing permafrost soils found AMGs associated with SOM degradation and central C metabolism, suggesting that viruses can play a direct role in augmenting the soil C cycle^92^. There were 773 total putative AMGs detected in 445 vMAGs within the FiRE database, including 138 CAZymes targeting diverse substrates such as cellulose, chitin, pectin, and xylan (**Supplementary data 6**). Additionally, the AMGs included 105 genes related to growth (e.g., ribosomal proteins, ribonucleoside-diphosphate reductase), 21 central carbon metabolism genes, and 21 genes encoding peptidases. Over 100 of these genes – including some related to SOM and necromass processing (e.g., glycoside hydrolases, polysaccharide lyases) and cell growth (pyrimidine ribonucleotide biosynthesis) – could be linked to all 8 of the featured High O Actinobacteria MAGs. Furthermore, transcripts mapped to 14 of this subset of AMGs, suggesting that prophage are manipulating SOM degradation metabolisms and potential cell growth rates of active bacterial MAGs in High O samples.

### Fungal genomes are active across burn conditions

Two fungal Ascomycota MAGs from taxa previously shown to be pyrophilous, *Leotiomycetes* (R113-184) and *Coniochaeata ligniaria* (R110-5)^68,94–96^, were reconstructed from the metagenomic data. These taxa were prominently represented in our ITS amplicon data; the *Leotiomycetes* class increased in relative abundance by ∼215% between control and High O samples (14% to 45%) and the *Coniochaeta* genus relative abundance increased from 0.003% to 1% from control to High A soils.

Complementing observations from pyrophilous bacterial MAGs, the fungal MAGs encoded and expressed genes for degrading aromatic compounds, including both the upper (peripheral) pathways for diverse aromatic substrates and lower pathways targeting intermediates catechol or protocatechuate. Each MAG expressed genes for degrading salicylate (salicylate hydroxylase), phenol (phenol 2-monooxygenase), and catechol (catechol 1,2-dioxygenase), and expression of all three genes increased from low to high severity in both soil horizons. The MAGs also encoded laccases, which are enriched in pyrophilous fungal genomes^97^ and act on aromatic substrates^98^. The *Coniochaeta* MAG additionally encoded hydrophobic surface binding proteins (*hsbA*; PF12296) which may facilitate the degradation of fire-derived hydrophobic compounds and be critical to soil recovery^97^. To directly compare the fungal and bacterial contribution to catechol degradation, we compared the number of normalized transcriptomic reads recruited to the gene encoding catechol 1,2-dioxygenase, *catA*. In High O samples, the fungal MAGs generated more than twice the number of transcripts per gene compared to bacterial MAGs, indicating the significant role that fungi likely play in aromatic DOM degradation in burned soils. Both fungal MAGs can also potentially degrade necromass from lysed heat-sensitive taxa through the expression of diverse peptidases (**Figure S12**), with increased expression from low to high fire severity in both organic and mineral horizons (∼40.4% and 235%, respectively).

### Ecosystem implications

We observed short-term changes in microbiome composition and function that likely alter biogeochemical cycling and initial post-fire vegetation recovery. Both bacterial and fungal community diversity decreased with burn severity (**Figure 2E, F**); a decrease in soil microbiome diversity can influence ecosystem function^99^. Despite the key role that bacterial N-fixing bacteria and mycorrhizal fungi play in generating plant available N pools^14^ (up to 80% of N acquired by plants in boreal and temperate forests), we found no expression of the functional gene catalyzing N fixation (*nifH*). Further, despite the three-fold increase in soil mineral soil ammonium observed in this study (**Supplementary data 1**) and others^100^, the gene that catalyzes ammonia oxidation (*amoA*) was absent from the MAG database. These observations mirror trends reported by other studies; short-term, post-fire decreases in the abundances of N-fixation and ammonia-oxidization genes have been noted in a mixed conifer forest^101^. Further highlighting potential reductions in metabolic function within the post-fire community, we also note that MAGs active in High O samples express fewer CAZymes that target a less diverse array of C substrates relative to High A MAGs (**Figure S6**), suggesting that certain C cycling processes may be absent in High O samples.

Ectomycorrhizal fungi (EMF) facilitate plant access to limiting nutrients and water in return for carbohydrates derived from photosynthesis^102^. We observed a 99% decrease in EMF relative abundance across the burn severity gradient (**Table S3**) from direct heat-induced or subsequent death of their plant hosts^18^, which likely has implications for obligate host plants such as *P. contorta*, the dominant tree species in these forests. For example, we found that *Cenoccum geophilum*, a common EMF symbiont of *P. contorta*^103,104^ was present in our unburned sites but absent after burning. Although we observed loss of EMF the year after severe wildfire, research in nearby forests indicates high (>90%) EMF colonization of pine roots within a decade of burning^105^.

Lastly, we show that wildfire and the post-fire soil chemical environment results in a microbiome that actively degrades aromatic compounds likely formed during fire (**Figure 5**). This finding supports recent studies suggesting that pyrogenic organic carbon is more labile than previously thought^106^, with implications for the modeling of C storage in wildfire impacted ecosystems. Further work is needed integrating multi-omics data from both field observations and laboratory experiments into ecosystem models to refine the quantification of C fluxes in post-fire ecosystems.

## Conclusion

Here we present FiRE-db, a publicly available, comprehensive genome database of pyrophilous microorganisms that provides functional context to observed shifts in soil microbiome structure following wildfire. Our results indicate that dominant microorganisms occupy available post-fire niche space through a combination of strategies, including heat tolerance, fast growth, and the utilization of substrates available post-fire. This ability to use aromatic DOM that is likely generated during wildfire is widespread throughout the bacterial and fungal community, with implications for assumptions regarding the residence time of pyrogenic carbon. Carbon processing within the system is also influenced by the presence and activity of abundant viruses that target key bacterial community members through viral predation and activity of AMGs. The measured patterns of community dynamics are both consistent across fires and trophic levels (i.e., bacteria, fungi, and viruses), offering opportunities to leverage these results for more effective management of other wildfire-disturbed environments.

## Acknowledgements

This work was supported through a USDA NIFA award (#2021-67019-34608) to MJW. FTICR MS analyses were performed under the Facilities Integrating Collaborations for User Science (FICUS) initiative and used resources at the Environmental Molecular Sciences Laboratory (proposal ID 49615), which is a DOE Office of Science User Facilities. This facility is sponsored by the Office of Biological and Environmental Research and operated under contract number DE-AC05-76RL01830. Metagenomic and metatranscriptomic sequencing was performed at the University of Colorado Cancer Center’s Genomics Shared Resource, which is supported by the Cancer Center Support Grant P30CA046934. This work utilized resources from the University of Colorado Boulder Research Computing Group, which is supported by the National Science Foundation (awards ACI-1532235 and ACI-1532236), the University of Colorado Boulder, and Colorado State University. The work conducted by the U.S. Department of Energy Joint Genome Institute, a DOE Office of Science User Facility, is supported by the Office of Science of the U.S. Department of Energy under Contract No. DE-AC02-05CH11231. We thank Fabiola Pulido-Chavez for help with processing the ITS amplicon sequencing data.

## Author contributions

ARN, CCR, and MJW designed the field sampling and downstream analyses. ARN, KKA, and MJW performed field sampling, while ARN and RAD performed laboratory sample processing. ARN and ABN led the microbial analyses, with assistance from SM, ASS, IG, and AS for fungal genomics. JBE and SEG assisted with viral interpretations, while HR, TB, RC, and RY contributed with high-resolution carbon measurements and analyses. TF performed bulk soil geochemistry measurements. ARN, CCR, and MJW wrote the manuscript, with assistance and input from all co-authors.

