## Supplementary text for "Playing with FiRE: A genome resolved view of the soil microbiome responses to high severity forest wildfire"

*Pyrophilous fungal taxa identified in burned samples*

The pyrophilous *Ascomycetes* increased in relative abundance by 118% between unburned and burned organic horizons (~34% to 75%; **Figure 1**); dominant genera included *Calyptrozyma* (~30% relative abundance in burned organic horizon), *Tricharina* (~13%), and *Geopyxis* (~7%). *Geopyxis* has been previously documented as a pyrophilous taxa and is identified as an endophyte, which would aid in their persistence through the fire event and proliferation thereafter[1,2]. The two most abundant *Ascomycete* orders in burned organic horizons were *Helotiales* (~31% relative abundance), which have been found in other post-fire soils[3,4] and lab pyrocosm experiments[5] and *Pezizales* (~28%), which is a common post-fire soil taxa[6–8] known for producing resistant structures like spores[9]. The second most dominant *Ascomycetes* family *Pyronemataceae* (~21% relative abundance in burned organic horizon) has been found to grow in soils heated in the laboratory to 70°C, to increase in forest soil samples post-fire, and to readily degrade aromatic compounds in the presence of pyrolyzed OM[6,10]. The *Ascomycotes* displayed a higher fire tolerance than the *Basidiomycota*, which had a decrease in relative abundance by ~58% between unburned and burned organic horizons, dropping from 50% to 21% relative abundance (**Figure 1**).

*Networks using Weighted Gene Correlation Network Analysis*

To determine how fire-induced changes in community richness translated into the complexity of potential interactions within the soil microbiome, we performed Weighted Gene Correlation Network Analysis[11] (WGCNA) on the 16S rRNA and ITS gene sequencing data. Across all O-horizon samples, we measured strongly decreasing numbers of nodes with increasing burn severity, and associated decreases in the edge-to-node ratio. Similar trends were apparent in the A-horizon samples, albeit with smaller changes between conditions (**Table S4**).

*Kendric mass-defect analyses*

Kendrick mass-defect (KMD) analysis with a base unit of $C_4H_2$ (equivalent to adding benzene to another benzene structure) provides further evidence of linkages between fire severity and increasing polyaromaticity (**Figure S13**). The series shown in Figure 4 reveal addition of a benzene ring between unburned and burned soils and an increase in relative abundance of the increasingly polyaromatic formulas (**Figure S13**)

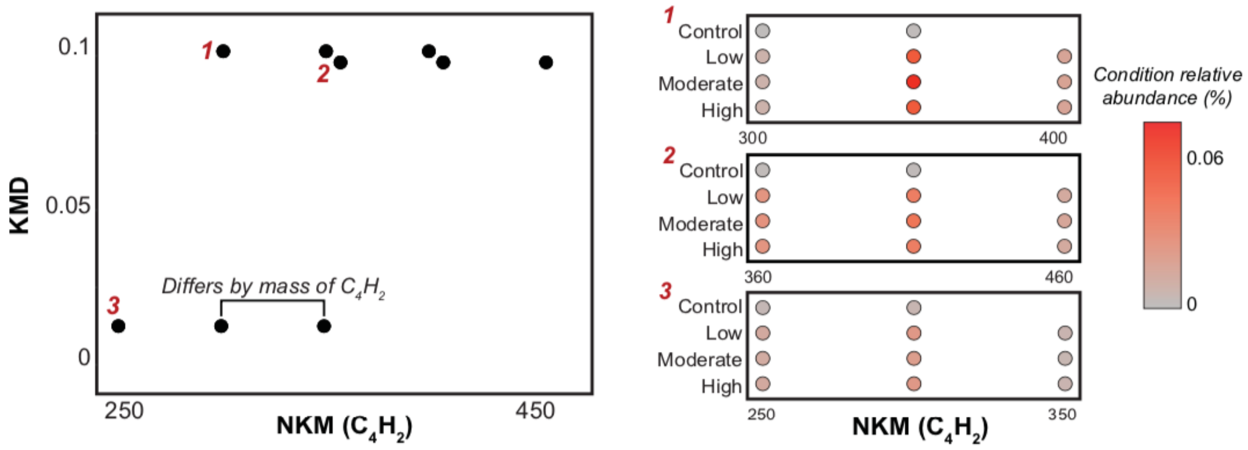


**Figure S13. (Left)** Example KMD series that differ by the mass of base unit $C_4H_2$ from samples that span the burn severity gradient. **(Right)** Each of the three series plotted across the burn severity gradient showing how the relative abundance differs and if a new formula is added with increasing severity.

*Genetic potential for processing polyaromatics*

We additionally looked for genes responsible for degrading naphthalene or removing a ring from a polycyclic aromatic ring (KEGG RM014; **Table S2**) and found that none of the MAGs encoded a naphthalene 1,2-dioxygenase (nahA), but 33 MAGs encoded 2-hydroxychromene-2-carboxylate isomerase (EC:5.99.1.4, nahD) and 10 encoded trans-o-hydroxybenzylidenepyruvate hydratase-aldolase (EC:4.1.2.45, nahE). These two genes were also expressed in the dataset, but only by two MAGs in the *Proteobacteria*.

*Broad functional shifts with burn severity*

To analyze broad shifts in functional potential and gene expression between low and high severity conditions within both soil horizons, we categorized genes using DRAM gene headers. There were 885,498 genes annotated with DRAM in the metagenomic dataset and 146,894 transcripts annotated with DRAM that had counts of at least 10 across all 12 samples. A DESeq2 analysis of the DRAM-annotated transcripts revealed 352 differentially expressed genes between Low O and High O soils and 26 between Low A and High A soils. The majority of differential expression in High O and High A was attributed to the Actinobacteria phyla (97.9% and 50% of differentially expressed genes, respectively). There were genera and family-level differences between the activity in both soil horizons, with *Actinobacteriota* genera *Blastococcus* (69.1%), *SCTD01* (10.4%) *Arthrobacter* (4.8%) and *Modestobacter* (2.7%) as the large players in High O and the family *Streptosporangiaceae* (35%) being the biggest player in High A. Outside of the *Actinobacteriota* phylum, the *Proteobacteria* genus *Palsa-1478* was responsible for 45% of differentially expressed genes in High A soils. Thus, the metatranscriptomics data additionally reflects the increased susceptibility of organic horizon soil to fire relative to mineral horizon soils.

There was a large change in broad functional potential between Low O and High O soils; of the 28 DRAM gene headers analyzed, 22 were significantly different in coverage between the severity conditions (**Figure S14**). Though there is a large shift in microbiome functional potential, the metatranscriptomics data reveals that there is less of a change in gene expression between Low O and High O conditions, exhibiting potential functional redundancy. Thirteen DRAM gene headers have differentially expressed genes, and 7 of these were genes only differentially expressed in high severity conditions (**Figure S14**). In total, there were 288 genes differentially expressed in High O and 64 genes differentially expressed in Low O. Reflecting the large role of *Actinobacteriota* in high severity-impacted soils mentioned above, the majority of differentially expressed genes in High O were from the *Actinobacteriota* phyla (97.9% of differentially expressed genes). Specifically, the *Actinobacteriota* genera *Blastococcus* (69.1%), *SCTD01* (10.4%) *Arthrobacter* (4.8%) and *Modestobacter* (2.7%)

Contrastingly, the microbiome of mineral soil horizons exhibited a much smaller shift in functional potential with only 4/28 DRAM gene headers having a significant change in coverage between Low A and High A (peptidase, information systems, CAZY, and CRISPR; **Figure S14**). There was also little change in gene expression; there were 6 DRAM headers (26 total genes) with differentially expressed genes, 20 that were differentially expressed in High A and 6 in Low A (**Figure S14**). The *Actinobacteriota* phyla was responsible for the largest proportion of differentially expressed genes following high-severity fire (50%) and the family *Streptosporangiaceae* and was responsible for 35% of the differentially expressed genes. The *Proteobacteria* genera *Palsa-1478* was also responsible for a large amount of differentially expressed genes (45%).

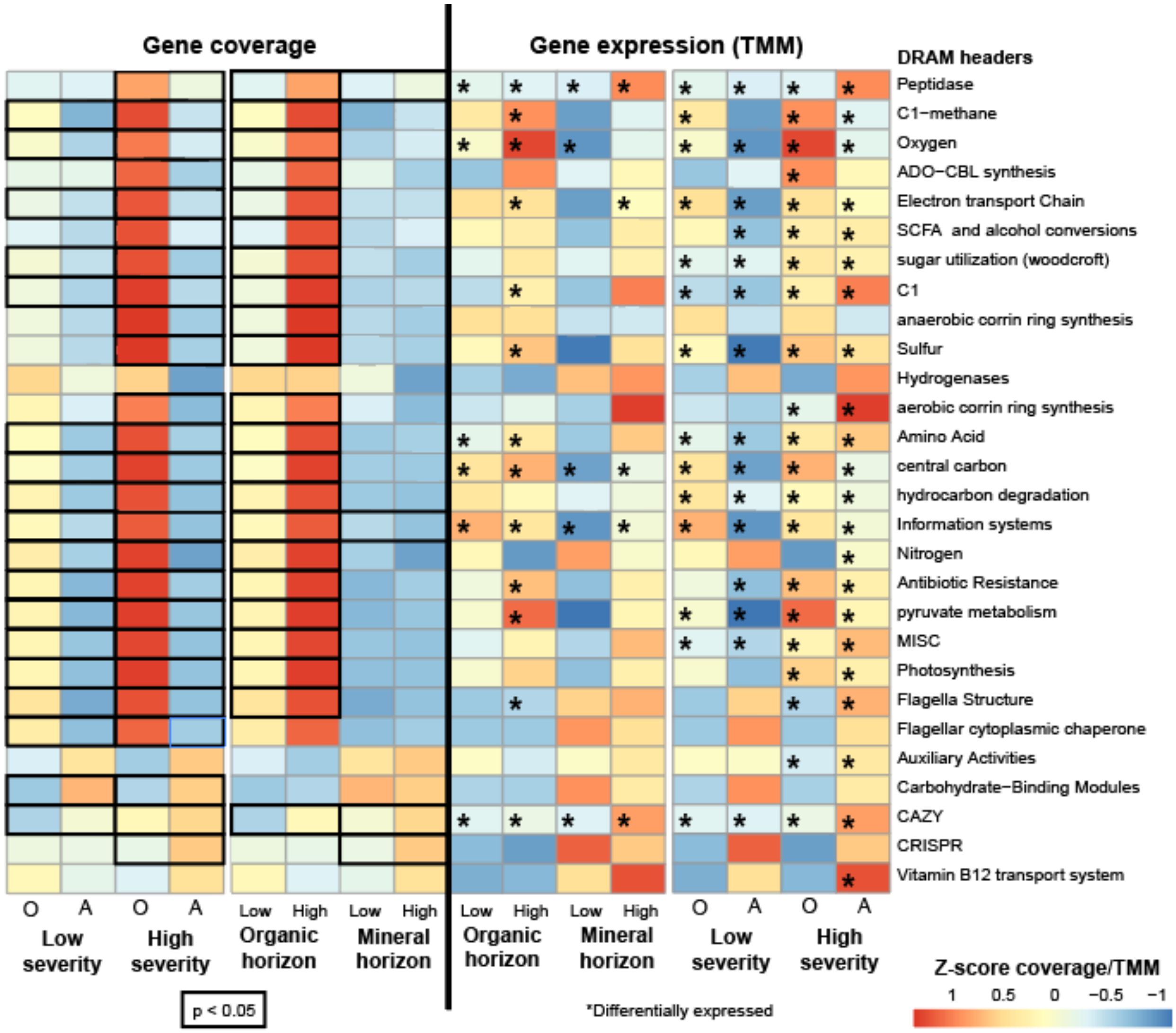


**Figure S14.** Overall trends between low and high severity organic and mineral horizons in the gene coverage (left) and gene expression (right) of the different DRAM gene headers. In the gene coverage heat maps, a bolded box indicates that the two conditions are significantly different from one another. In the gene expression data, stars indicate whether there are genes within that DRAM header that are differentially expressed in that condition relative to the other.

*Fungal MAGs encode secondary metabolite clusters*

The *Coniochaeta* MAG (R110-5) encoded 26 secondary metabolite clusters, including 12 PolyKetide Synthase (PKS) clusters and 4 Non-Ribosomal Peptide Synthetase (NRPS) clusters. This trait appears to be conserved across the *Coniochaeta* genera, as the other four *Coniochaeta* genomes deposited on JGI MycoCosm encode an average of ~36, ranging from 25-66. Both polyketide and non-ribosomal peptide secondary metabolites display a diverse array of biological uses and properties including antibiotic synthesis[12], siderophore production[13], and fungal developmen[14] (e.g., sporulation factors), and may give organisms a competitive edge, especially under stressful conditions. The other fungal MAG (R113-184; *Leotiomycetes*) may persist following wildfire due to an endophytic lifestyle; 36 *Leotiomycetes* ESV's were classified as endophytes by FUNGuild. Pyrophilous endophytic fungi have been found to occur survive fire events in small-scale refugia then, triggered by the fire, produce reproductive sporocarps[1].
