## Supplementary material for "Playing with FiRE: A genome resolved view of the soil microbiome responses to high severity forest wildfire": Figure S

**Supplementary figures**

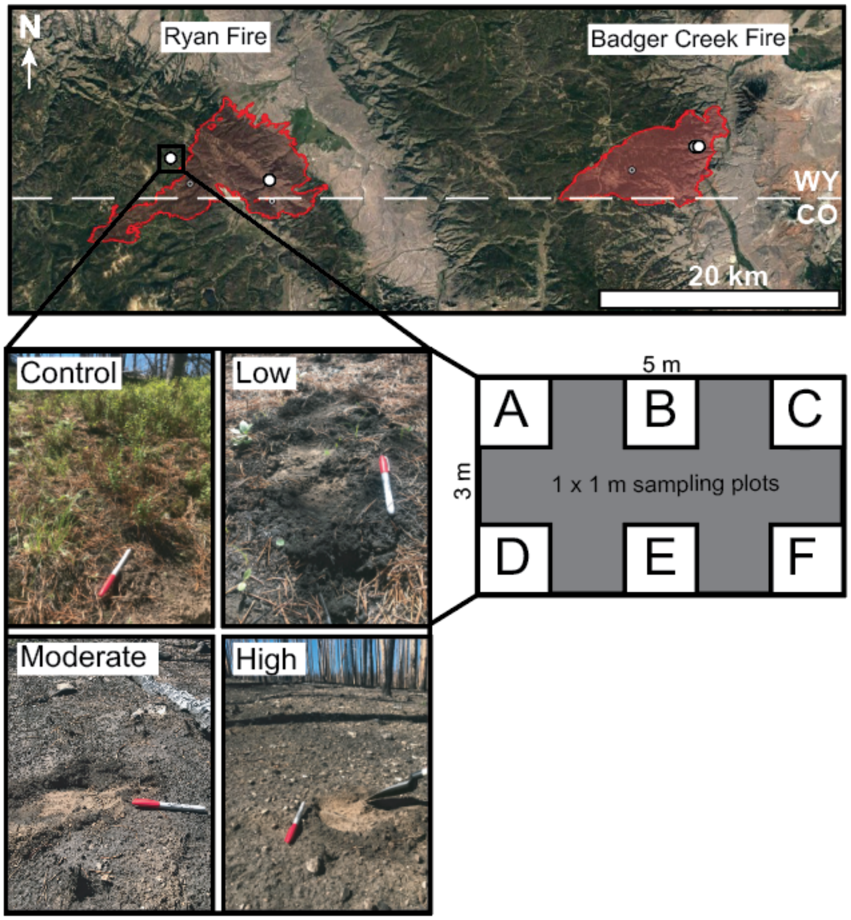


**Figure S1.** Schematic of sampling design. There were four replicate burn severity gradients (two at Ryan Fire and two at Badger Creek Fire); six subsamples were collected in each burn condition at each gradient. Burn severity classes were based on Parson et al. (2010).

| Sample | TMM high expression cutoff |
|---|---|
| **R85** | 1.68 |
| **R86** | 3.26 |
| **R89** | 1.47 |
| **R90** | 3.16 |
| **R93** | 1.68 |
| **R94** | 3.28 |
| **R109** | 1.68 |
| **R110** | 2.24 |
| **R113** | 1.68 |
| **R114** | 2.11 |
| **R117** | 1.61 |
| **R118** | 1.87 |

**Table S1.** TMM cutoff values for high expression analysis in each sample. Values correspond to the 20$^{th}$ percentile value TMM. Transcripts were highly expressed if the TMM was above this value for 2/3 samples in any one condition.

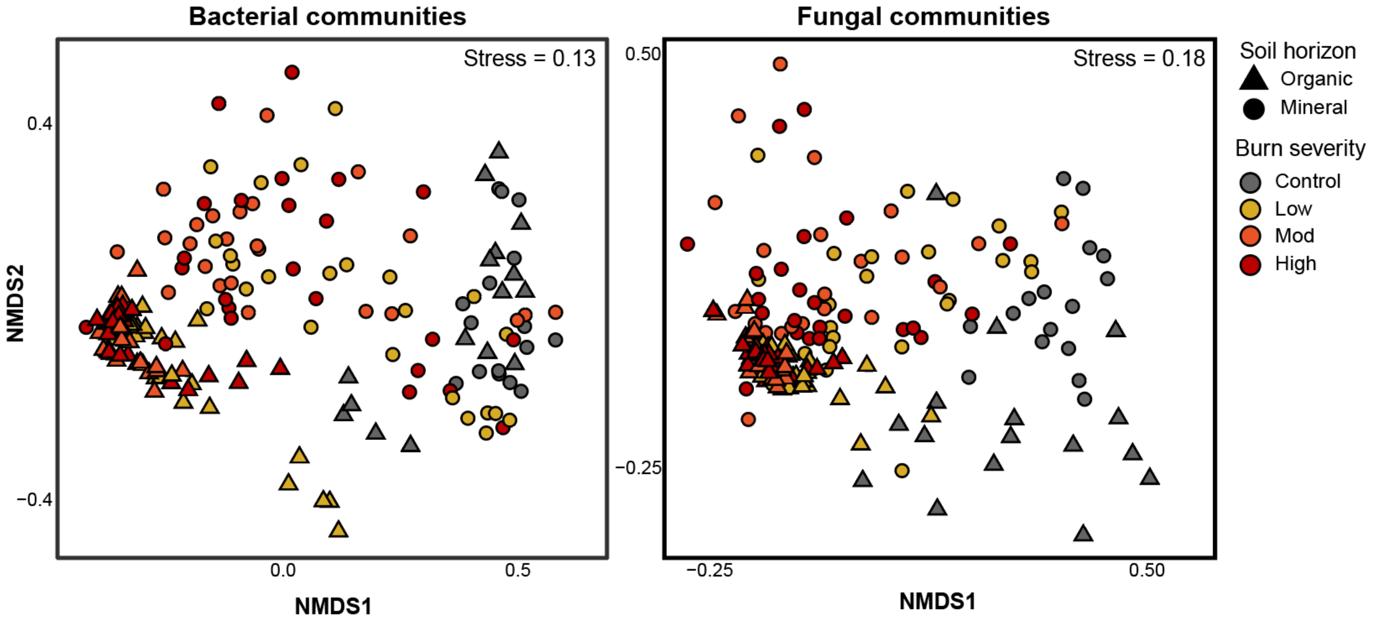


**Figure S2.** Non-metric multidimensional scaling (NMDS) ordination of all bacterial (left) and fungal (right) communities shaped by soil horizon and colored by burn severity.

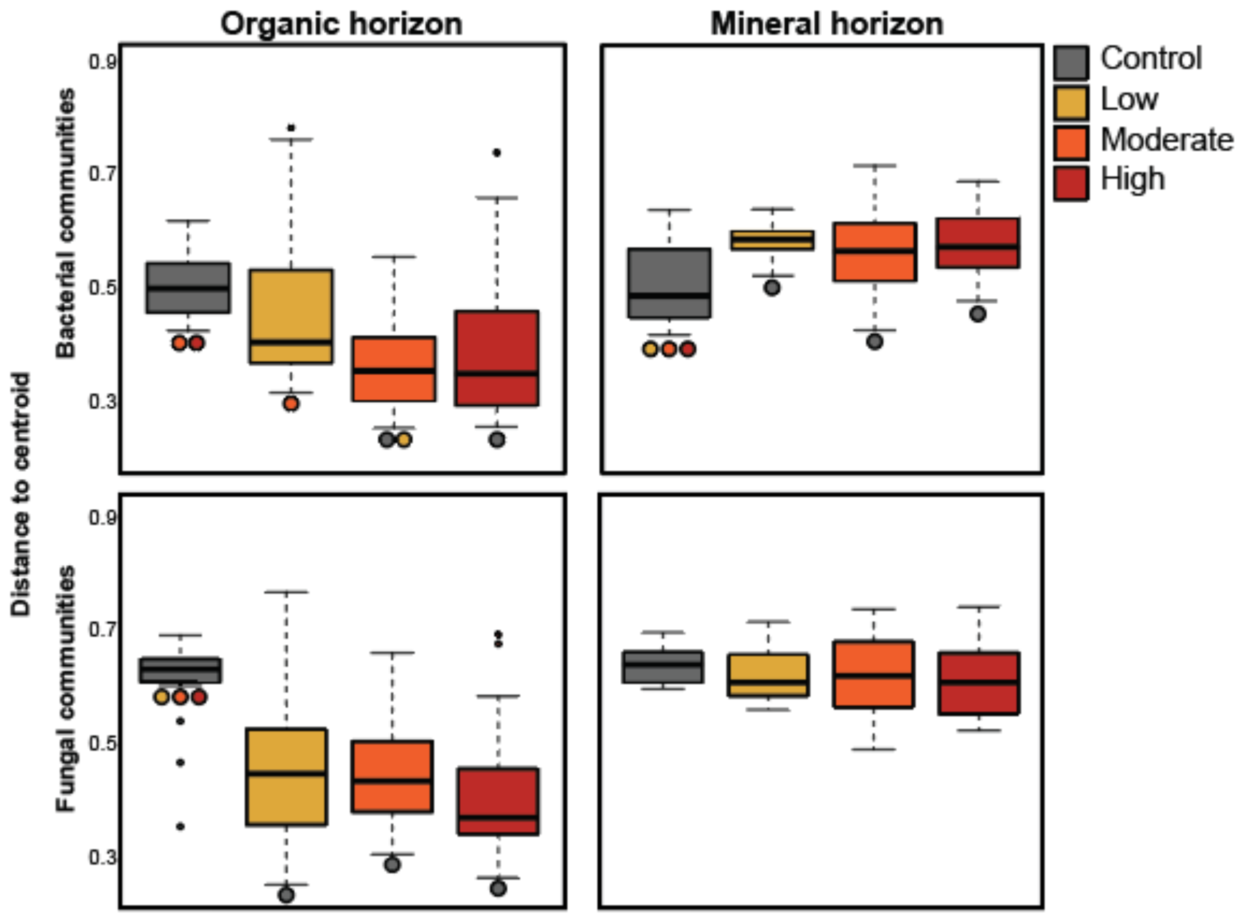


**Figure S3.** Distance to centroid calculations of the NMDS ordinations or O-horizon and A-horizon bacterial (top) and fungal (bottom) communities by burn severity. Colored points indicate significant differences ($p<0.05$) between conditions.

| Condition | Nodes | Edges | Edge:node | Modules | N samples |
|---|---|---|---|---|---|
| **Bacterial communities** | | | | | |
| **Control, organic** | 10426 | 3662534 | 351.3 | 16 | 16 |
| **Low, organic** | 6234 | 1194870 | 191.7 | 22 | 24 |
| **Moderate, organic** | 2442 | 196930 | 80.6 | 11 | 24 |
| **High, organic** | 2193 | 277073 | 126.3 | 11 | 24 |
| **Control, mineral** | 9452 | 2769191 | 292.9 | 16 | 16 |
| **Low, mineral** | 9132 | 1904471 | 208.5 | 21 | 24 |
| **Moderate, mineral** | 5916 | 1376628 | 232.7 | 18 | 24 |
| **High, mineral** | 6415 | 1151930 | 179.6 | 19 | 24 |
| **Fungal communities** | | | | | |
| **Control, organic** | 2491 | 271323 | 108.9 | 12 | 16 |
| **Low, organic** | 1548 | 126698 | 81.8 | 11 | 24 |
| **Moderate, organic** | 238 | 14366 | 60.3 | 2 | 24 |
| **High, organic** | 260 | 17026 | 65.5 | 2 | 24 |
| **Control, mineral** | 1569 | 145649 | 92.8 | 9 | 16 |
| **Low, mineral** | 681 | 48389 | 71.1 | 5 | 24 |
| **Moderate, mineral** | 242 | 14558 | 60.1 | 3 | 24 |
| **High, mineral** | 374 | 23626 | 63.1 | 3 | 24 |

**Table S2.** Characteristics of WGCNA networks created from 16S rRNA gene sequencing data from each condition.

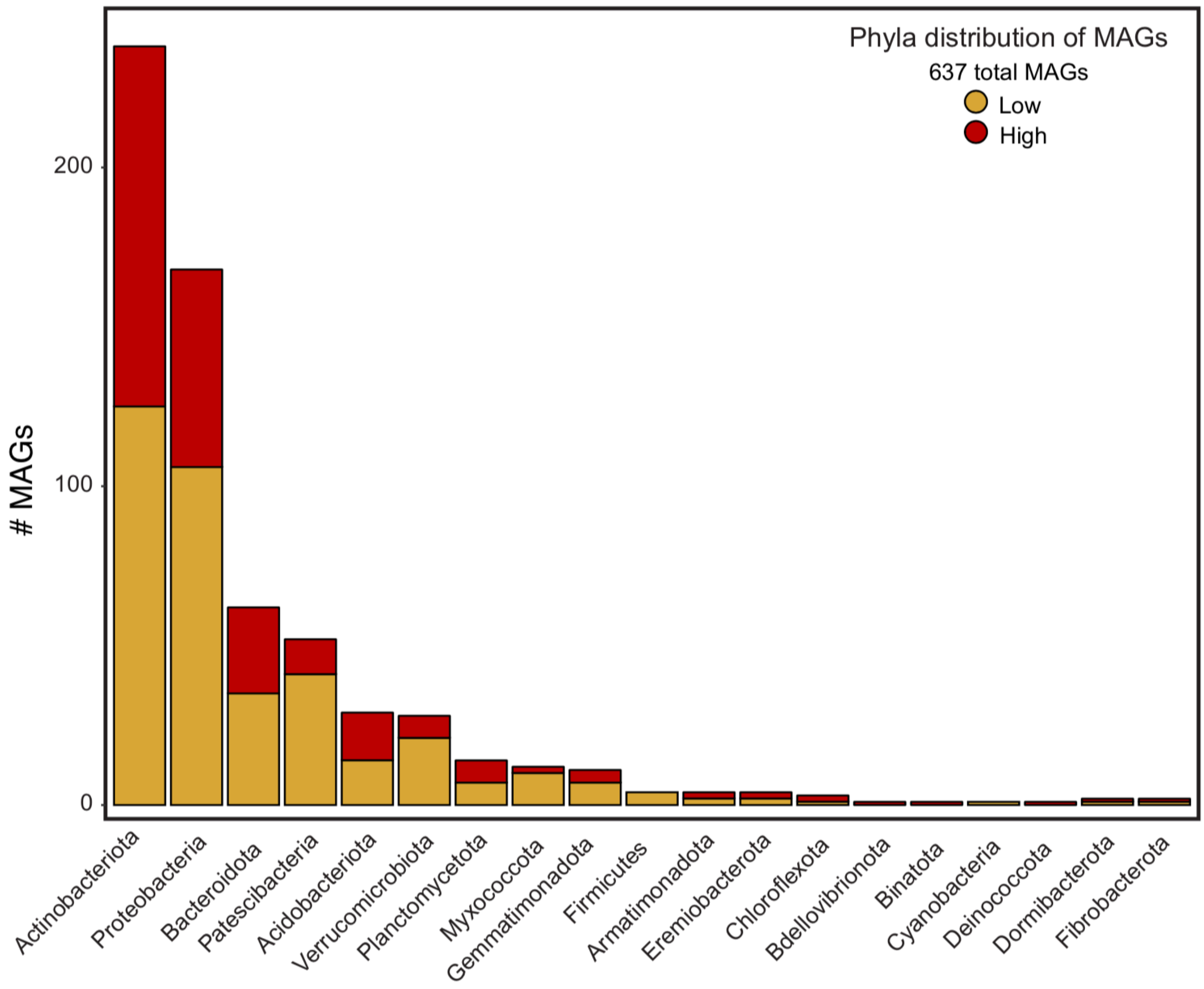


**Figure S4.** Phyla distribution of the 637 medium- and high-quality MAGs (> 50% completion, <10% contamination) from burned O- and A-horizon soils.

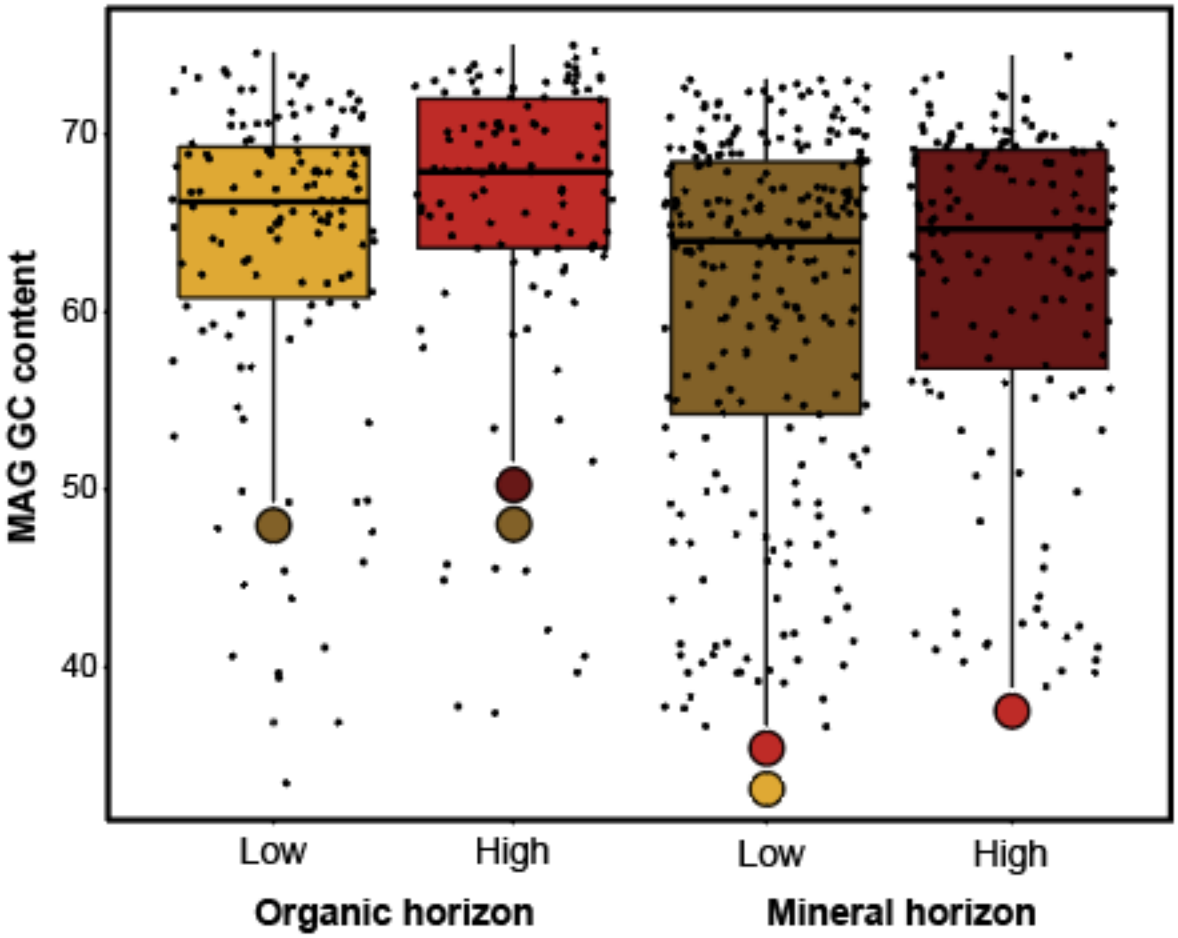


**Figure S5.** GC content of MAGs reconstructed from each condition. Colored points indicate significance (p<0.05) between conditions.

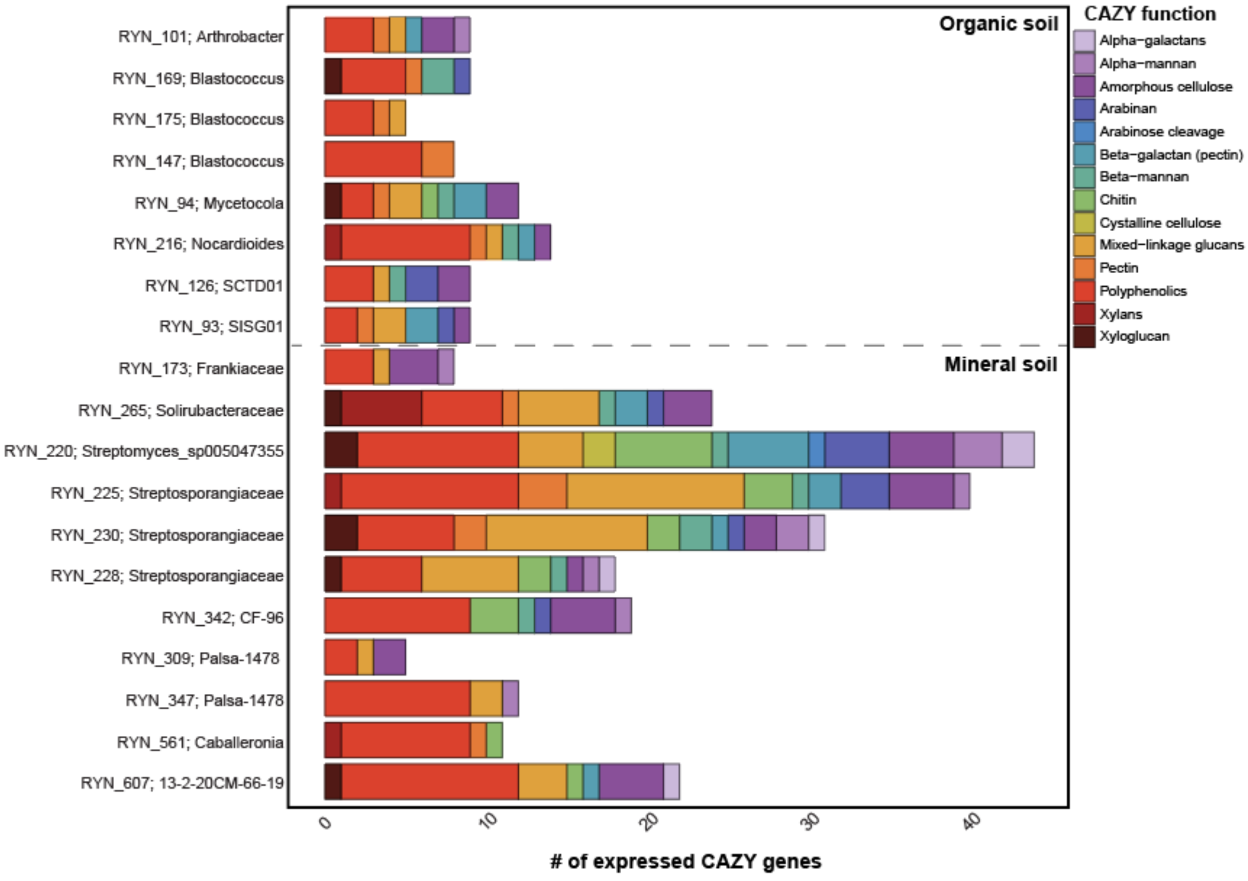


**Figure S6.** Functional diversity of expressed CAZYmes of the MAGs of interest from High O and High A soils.

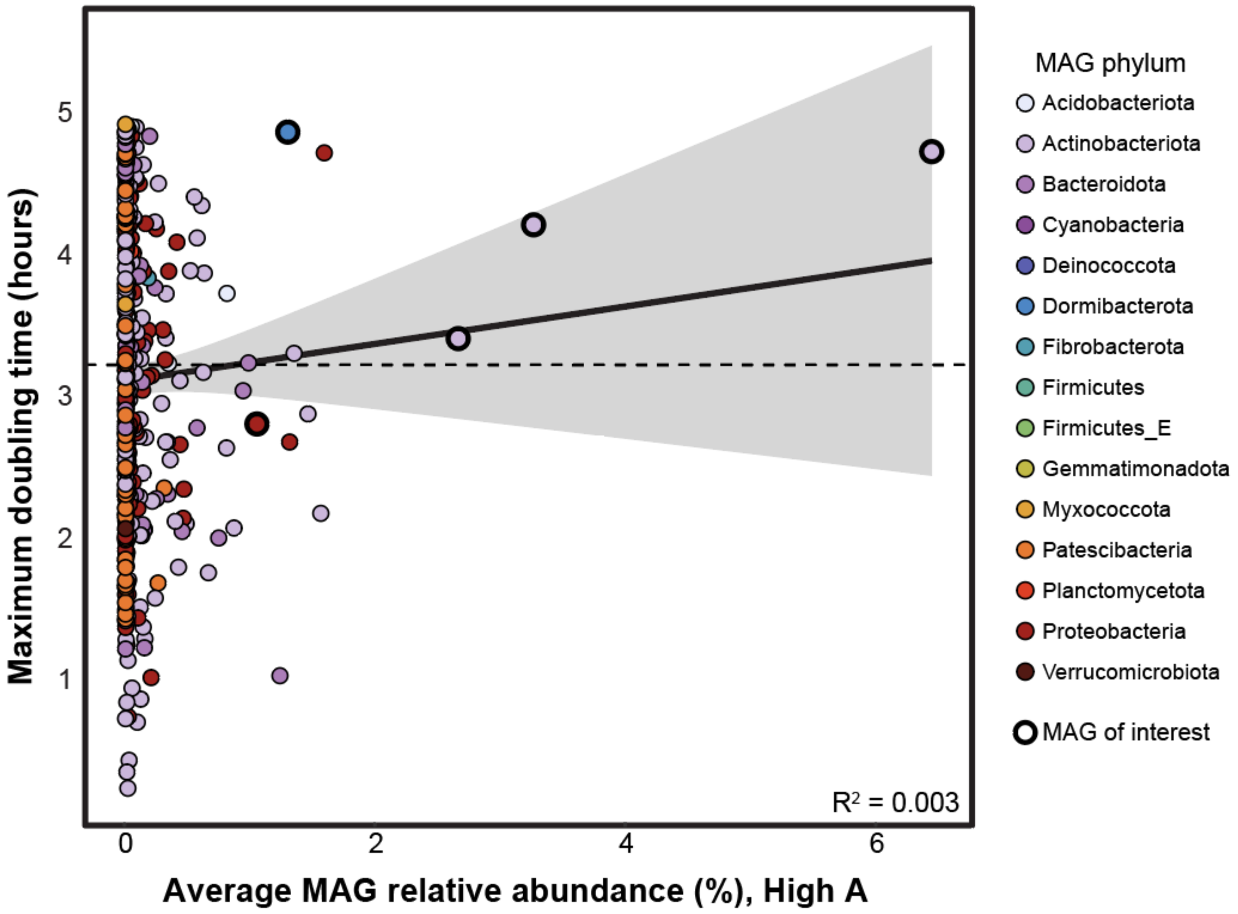


**Figure S7.** The average relative abundance of the MAG in High A soils plotted against maximum doubling time. MAGs of interest are bolded.

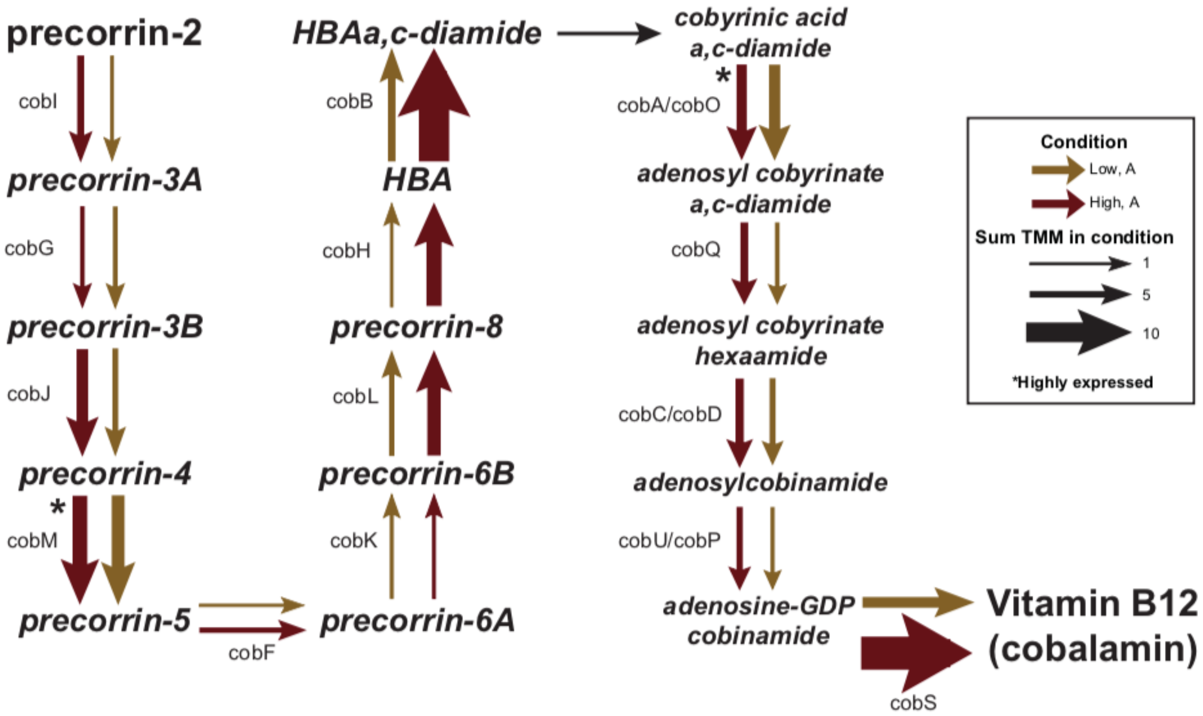


**Figure S8.** The aerobic cobalamin biosynthesis pathway (adapted from Doxey et al., 2015 and Lu et al., 2020) with arrows indicating the summed TMM in the colored condition.

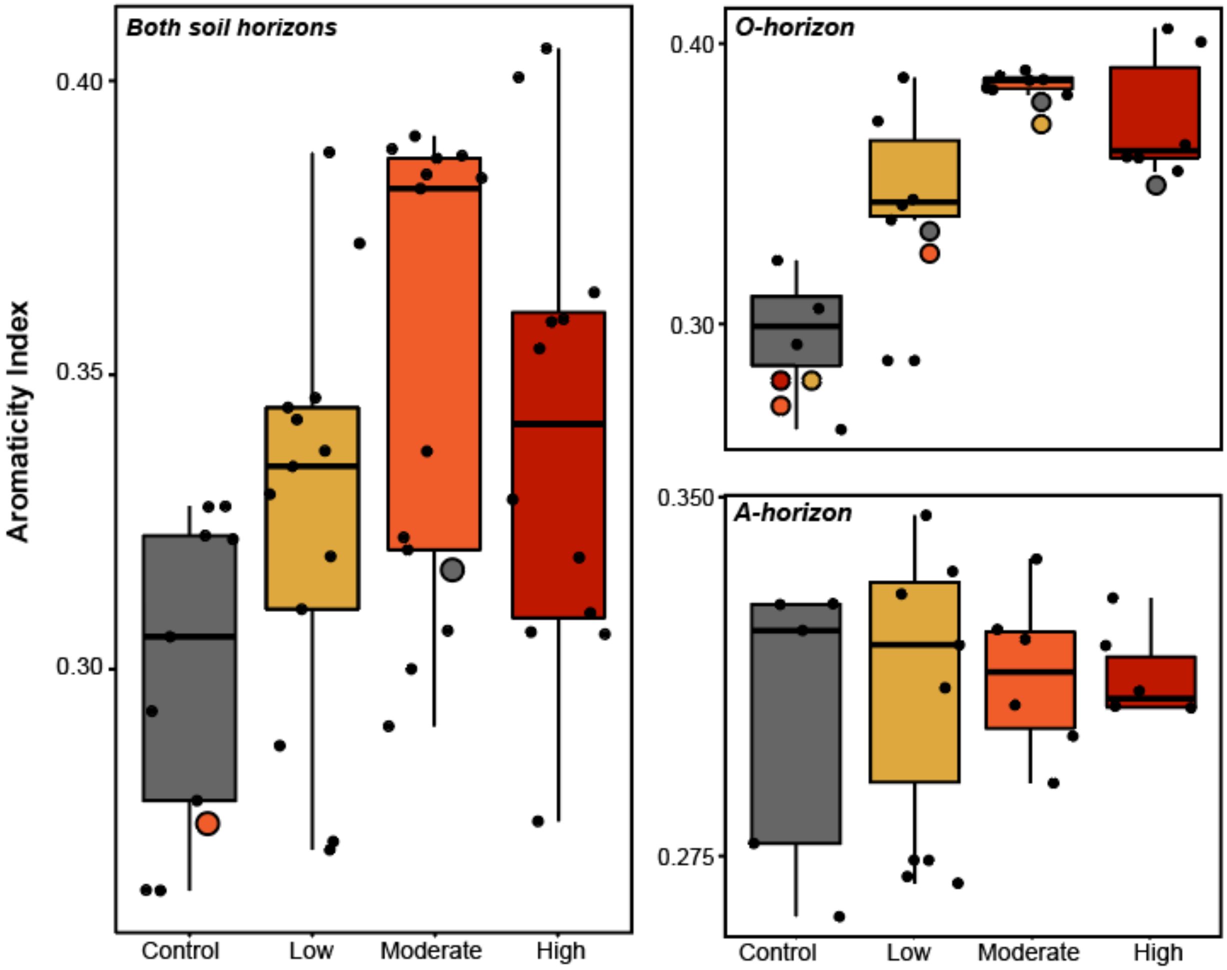


**Figure S9.** Aromaticity index of DOM pools against burn severity conditions and soil horizons. Colored circles indicate significance between the indicated conditions.

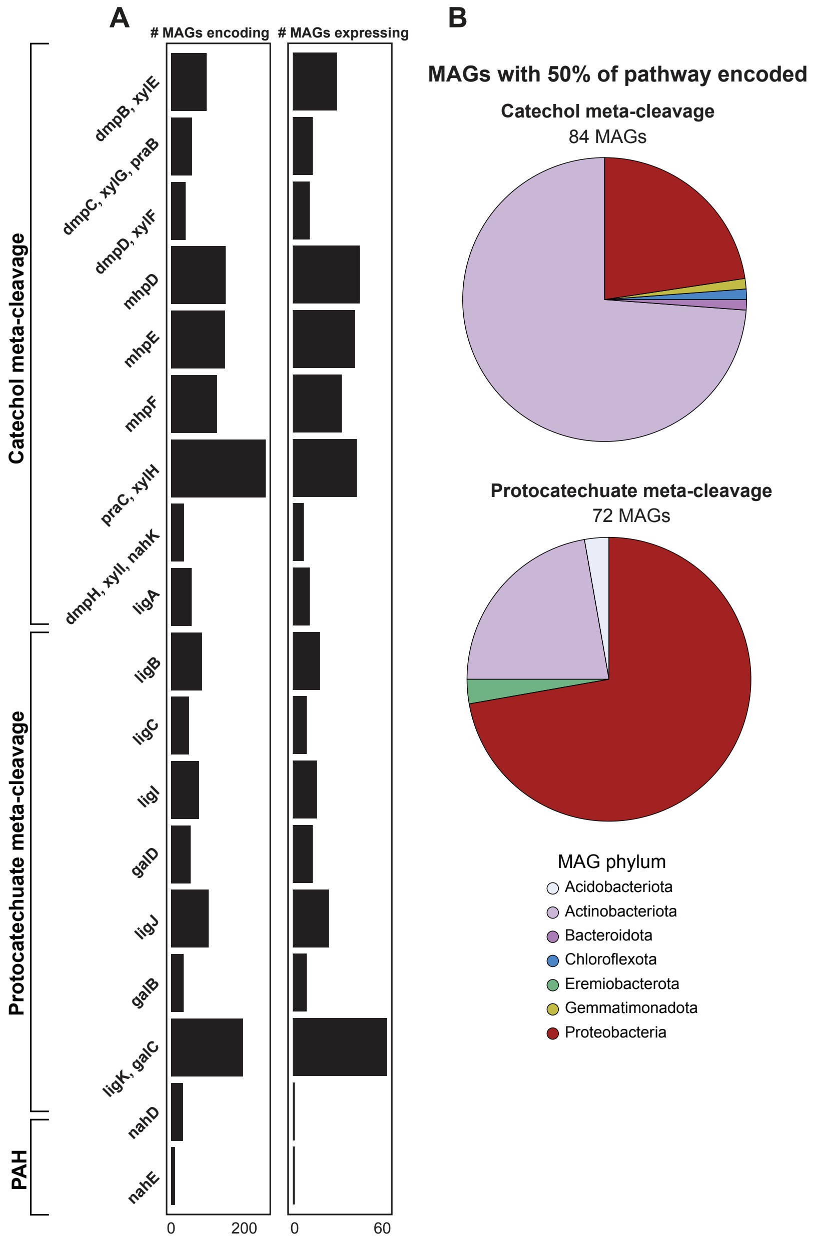


**Figure S10. (A)** Number of MAGs encoding and expressing each gene of the catechol and protocatechuate meta-cleavage pathways. **(B)** Phyla distribution of MAGs encoding 50% of either pathway.

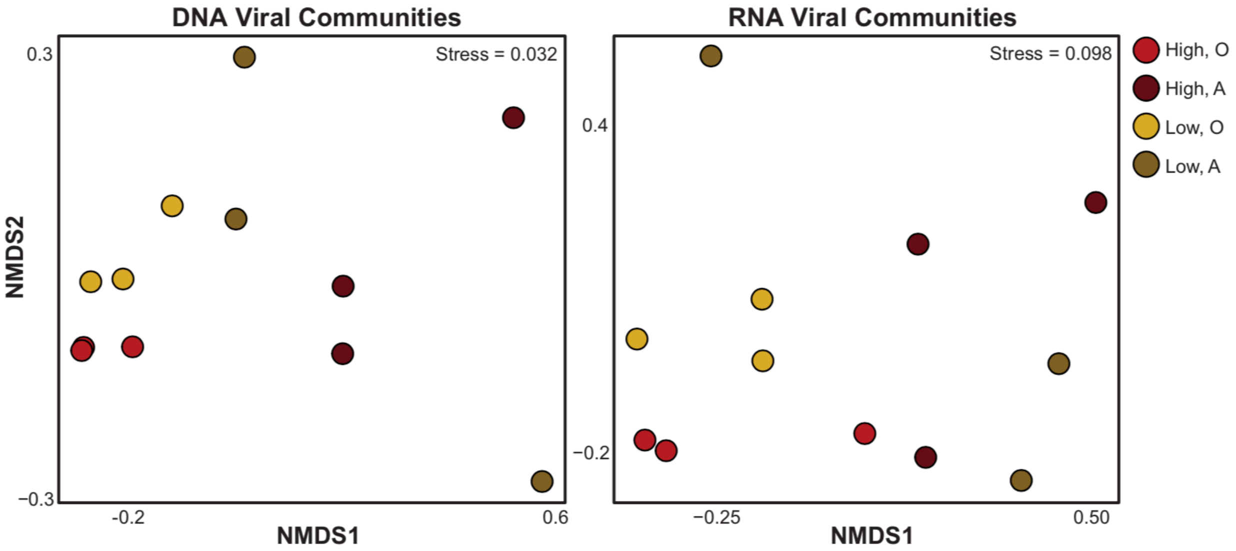


**Figure S11.** NMDS of DNA (left) and RNA (right) vMAG abundance across all four conditions.

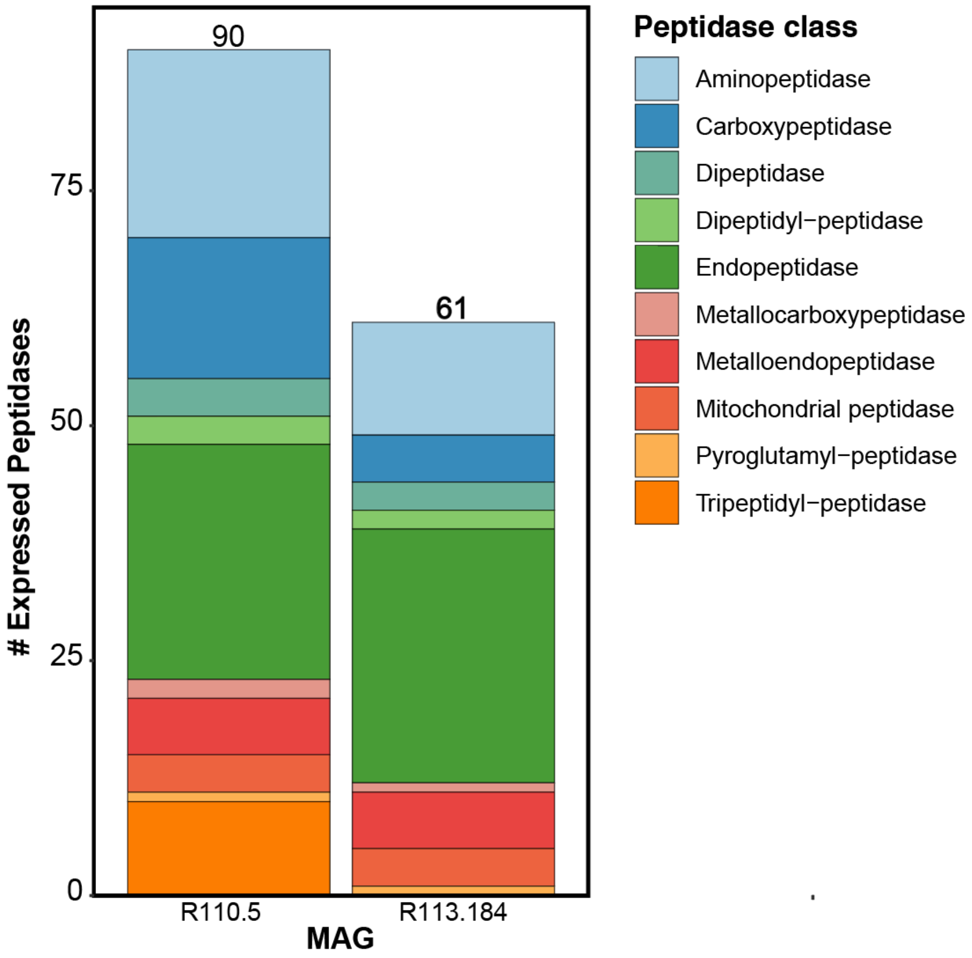


**Figure S12**. The diversity of peptidases expressed by each fungal MAG across all 4 conditions.

| Ecological guild | # ESVs | % Change with Low | % Change with Mod | % Change with High |
|---|---|---|---|---|
| *Arbuscular Mycorrhizal* | 17 | -100 | -96.65 | -100 |
| *Ectomycorrhizal* | 321 | -99.71 | -99.93 | -99.92 |
| *Endophyte* | 39 | -94.26 | -95.96 | -99.91 |
| *Epiphyte* | 31 | -81.06 | -99.34 | -98.78 |
| *Saprotroph* | 1624 | 228.87 | 260.75 | 281.42 |

**Table S3.** Percent change in relative abundance from control to low, moderate, and high severity in O-horizon soils of all ESVs assigned to each ecological guild by FUNguild.
